# OligoFragger: spectrum graph-guided analysis of modified RNA oligonucleotides by mass spectrometry

**DOI:** 10.64898/2026.11.09.765503

**Authors:** Jiechen Shen, Daniel A. Polasky, Fengchao Yu, Kaley M. Simcox, Bojing Zhu, Kristin S. Koutmou, Alexey I. Nesvizhskii

## Abstract

RNA modifications are widespread modulators of RNA function, but comprehensive identification remains technically challenging. This is especially true for densely modified RNAs such as tRNAs, where multiple modified residues and many possible chemistries occur within short sequence contexts. Tandem mass spectrometry provides direct chemical and positional evidence for modified oligonucleotides, but current database-search strategies struggle when dozens of candidate modifications must be considered simultaneously. Here, we present OligoFragger, a spectrum graph-guided computational tool that uses fragment evidence to constrain modified-oligonucleotide searches before full database scoring, enabling rapid and robust oligonucleotide identification. Using controlled RNA samples and tRNA datasets from *E. coli* and human cells, we demonstrate that OligoFragger increases identifications under stringent FDR control while recovering known modification patterns and supporting label-free quantification. This strategy enables sensitive, FDR-controlled identification and label-free quantification of modified RNA oligonucleotides in complex epitranscriptomic search spaces.

## INTRODUCTION

RNA modifications form a diverse and functionally important layer of gene regulation, collectively referred to as the "epitranscriptome"^1,2^. More than 170 chemically distinct modifications have been identified across all domains of life, with transfer RNAs (tRNAs) being the most heavily modified RNA species^1,3,4^. These modifications modulate codon recognition, translational fidelity, tRNA stability, and cellular stress responses^5,6^. Ribosomal RNAs (rRNAs) also carry numerous modifications that are critical for ribosome assembly and function^7^. More recently, modifications in messenger RNAs (mRNAs) and long non-coding RNAs (lncRNAs) have been reported to influence splicing, translation efficiency, and RNA degradation^8,9^. RNA modifications are also increasingly linked to disease mechanisms and the design of RNAs, including modified mRNA vaccines and synthetic small RNA therapeutics^10^. Comprehensive mapping of RNA modifications is therefore essential for understanding their biological functions, clinical relevance, and roles in engineered RNA molecules.

Several approaches have been developed to detect RNA modifications, each with its own strengths and limitations^11,12^. Sequencing-based approaches typically convert RNA into complementary DNA (cDNA) by reverse transcription (RT) and infer modifications from RT-induced signatures such as misincorporations, truncations, or stops. These approaches can be sensitive and applied across the transcriptome, but many modifications are detected indirectly, and the readout depends on modification chemistry, sequence context, and the behavior of the reverse transcriptase^11,12^. Nanopore direct RNA sequencing provides a complementary approach: native RNA passes through a protein nanopore, and changes in the ionic current can, in principle, reveal modified residues without cDNA conversion. In practice, however, modification calling depends on the models and training data used. Recent benchmarking of RT and nanopore RNA-modification tools has highlighted challenges in precision, quantification, and biological validation^13–15^.

Mass spectrometry (MS) offers an orthogonal, chemically direct way to identify and quantify RNA modifications, and is particularly attractive for heavily modified RNA species such as tRNAs^16–19^. Two strategies are commonly used: nucleoside analysis and oligonucleotide (bottom-up) analysis. In nucleoside analysis, RNA is digested exhaustively into single nucleosides, which are separated and detected by LC-MS/MS to provide a global inventory of modified residues. While highly sensitive for detecting and quantifying individual modified nucleosides, this approach loses positional information and cannot assign modifications to specific sites in the original RNA sequence. In contrast, the bottom-up oligonucleotide workflow preserves this positional information: RNA is enzymatically digested with sequence-specific ribonucleases (e.g., RNase T1, which cleaves after guanosine, or RNase A, which cleaves after pyrimidines) to generate oligonucleotide fragments amenable to LC-MS/MS analysis^18,20^. These oligonucleotides are then fragmented in the mass spectrometer, producing characteristic ion series (a, b, c, d from the 5’ terminus and w, x, y, z from the 3’ terminus) that enable sequence determination and modification site localization^21–23^. RNA oligonucleotide MS/MS also presents mass ambiguities: the same nucleotide sequence can be observed with different terminal phosphate or dehydrated-phosphate states, changing the precursor mass without changing the sequence itself^24,25^.

Despite the maturity of these experimental workflows, computational tools for RNA oligonucleotide MS/MS remains less developed than that for proteomics^26,27^. Existing tools, including Ariadne^28^, RoboOligo^29^, RNAModMapper/RAMM^30^, NucleicAcidSearchEngine (NASE)^31^, BioPharma Finder, and Pytheas^32^, have made important contributions to RNA sequence identification, interpretation of modified-oligonucleotide, target-decoy validation, visualization, and automated analysis. Nevertheless, most of these workflows remain sequence-first: candidate oligonucleotides are generated by in silico digestion, modified variants and terminal mass states are considered during database construction or matching, and theoretical spectra are compared against experimental MS/MS data. This strategy is effective for focused searches, but it becomes limiting for highly modified RNAs such as tRNAs. For example, a reference catalog of *E. coli* tRNAs includes 28 modification types, with distributions that depend on nucleotide position and tRNA isoacceptor^33^. Searching dozens of variable modifications while allowing multiple modified sites within each oligonucleotide creates a large combinatorial search space. Searching more modifications and terminal states expands the target and decoy candidate spaces, increasing the number of plausible but incorrect matches that compete for each spectrum. Under target-decoy FDR control, this added competition can force more stringent score cutoffs and reduce the number of accepted identifications, particularly when fragment evidence is incomplete or noisy. These challenges motivate search strategies that use fragment-level evidence earlier in the workflow to constrain candidate compositions, terminal states and modification possibilities before full scoring.

Here, we introduce OligoFragger, a computational method and software for identifying and quantifying modified RNA oligonucleotides by mass spectrometry. OligoFragger uses fragment evidence to guide the search for modified sequences. This approach adapts spectrum-graph and sequence-tag strategies that made searches for many modifications tractable in proteomics^34–36^ to RNA oligonucleotides. For each MS/MS spectrum, OligoFragger constructs a directed graph from the deisotoped peaks, connecting peaks whose mass differences correspond to individual nucleotides, including modified nucleotides. Paths through the graph provide sequence and modification information that is used to select candidates from an unmodified sequence database. This avoids enumerating all possible modified sequence variants before searching and limits full scoring to candidates supported by the observed fragments. Candidate sequences are checked against the precursor mass, accounting for alternative phosphate and water-loss states, and scored using the number and normalized intensities of matched fragments. Low-mass diagnostic ions can further restrict the modifications considered in each spectrum. OligoFragger also flags modification assignments with weak or conflicting fragment evidence for further inspection. Statistical confidence in oligonucleotide-spectrum matches is assessed using target-decoy competition, with FDR estimation performed separately for each file. MS1-based label-free quantification is provided through IonQuant^37^. We demonstrate OligoFragger’s performance through controlled validation experiments, comprehensive *E. coli* tRNA profiling, benchmarking on human tRNA data and reproducible label-free quantification.

## RESULTS

### OligoFragger workflow

The OligoFragger workflow was designed to search modified RNA oligonucleotide spectra without first expanding the database into all possible modified sequence variants (Fig. 1a). During database preparation, input RNA sequences are digested in silico and paired with shuffled decoys for FDR estimation, but the target+decoy database remains an unmodified sequence space. Searchable modifications are introduced later, at the spectrum-graph stage. This avoids combinatorial expansion of modified sequence databases and distinguishes OligoFragger from sequence-enumeration approaches such as NucleicAcidSearchEngine (NASE).

**Figure 1.**
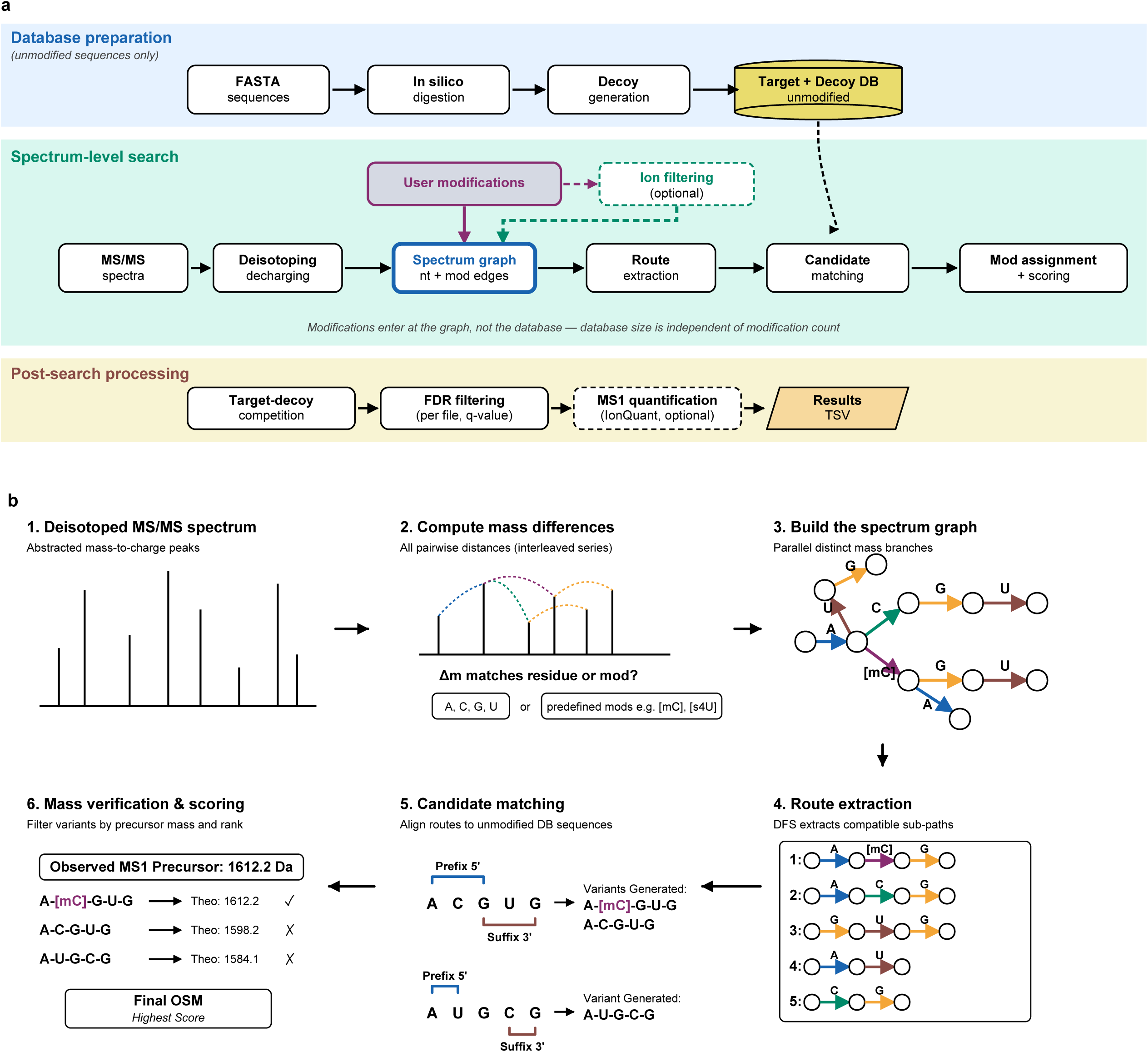
OligoFragger workflow for RNA oligonucleotide MS/MS analysis. **(a)** Three-phase pipeline overview. Database preparation generates in silico digestion products and shuffled decoys from input FASTA sequences while keeping the target+decoy database in unmodified sequence space. Spectrum-level search introduces user-selected modifications during spectrum graph construction, where modified residues contribute possible graph edges rather than pre-enumerated database entries. Post-search processing applies target-decoy competition, FDR filtering, suspicious-modification annotation, optional MS1 quantification, and results export. **(b)** Schematic of the spectrum-level search for a single spectrum. Deisotoped MS/MS peaks are converted into graph nodes, pairwise mass differences are matched to canonical or modified residue masses, and matched differences define directed graph edges. Routes through the graph provide spectrum-supported sequence tags; when a route uses a modified edge, that modification is carried forward as part of the candidate evidence. These routes are matched against unmodified database candidates to generate compatible modification-aware variants, which are then verified by precursor mass and ranked by route-evidence scoring. The highest-scoring validated target or decoy assignment becomes the oligonucleotide-spectrum match.

Each MS/MS spectrum is then searched in a spectrum-first manner (Fig. 1b). Deisotoped peaks are represented as graph nodes, and edges are drawn when the mass difference between two peaks matches a canonical ribonucleotide or a user-selected modified residue. In this representation, adding a modification to the search adds possible edge labels to the spectrum graph rather than new entries to the target-decoy database. Optional diagnostic ion filtering can further restrict the modification set to residues with spectrum-level low-mass evidence before candidate scoring.

Routes through the graph provide spectrum-supported sequence tags that already carry modification identity when a modified edge is used. These routes are matched to unmodified database candidates, and compatible routes generate a focused set of modification-aware sequence variants for each candidate. The variants are then checked against the observed precursor mass and scored using matched route evidence from RNA backbone cleavage ions. Because RNA digestion and ion activation can produce the same oligonucleotide sequence with different terminal phosphate or dehydrated-phosphate states, OligoFragger also evaluates terminal mass offsets during precursor validation and uses matched terminal-ion evidence to restrict the appropriate state when possible (Supplementary Note 1). The highest-scoring validated target or decoy assignment is retained as the oligonucleotide-spectrum match (OSM) for that spectrum.

After spectrum-level searching, OligoFragger applies target-decoy competition and calculates q-values separately for each input file. Accepted matches are reported with matched-fragment evidence, route support, suspicious-modification annotations, and, when MS1 data are available, IonQuant-based label-free intensities. OligoFragger uses spectrum evidence to flag modification assignments that are likely to be false; these flags are reported for downstream interpretation rather than used to automatically remove identifications.

### Validation with controlled samples

#### Reanalysis of NASE lncRNA NME1 with NCL1 methyltransferase treatment

To validate OligoFragger’s ability to detect methylation events, we reanalyzed the NASE dataset generated from a 340-nt, in vitro-transcribed yeast NME1 RNA, with and without treatment by the 5-methylcytidine (m^5^C) methyltransferase NCL1^31^. NCL1 is known to catalyze the formation of m^5^C on RNA substrates. NME1 was searched with a focused methylated-base panel containing mA, mC, mU and mG(denoting monomethylated nucleosides without specifying the chemical position of methylation), allowing methylcytidine assignments to compete with other common methylated-base assignments. In the untreated control runs, OligoFragger identified 1,648 OSMs at 1% FDR, of which only two carried non-suspicious modification assignments. Both were mC-containing OSMs (0.12% of accepted OSMs), showing that methylcytidine assignments remained rare in the absence of enzymatic treatment (Fig. 2a).

**Figure 2.**
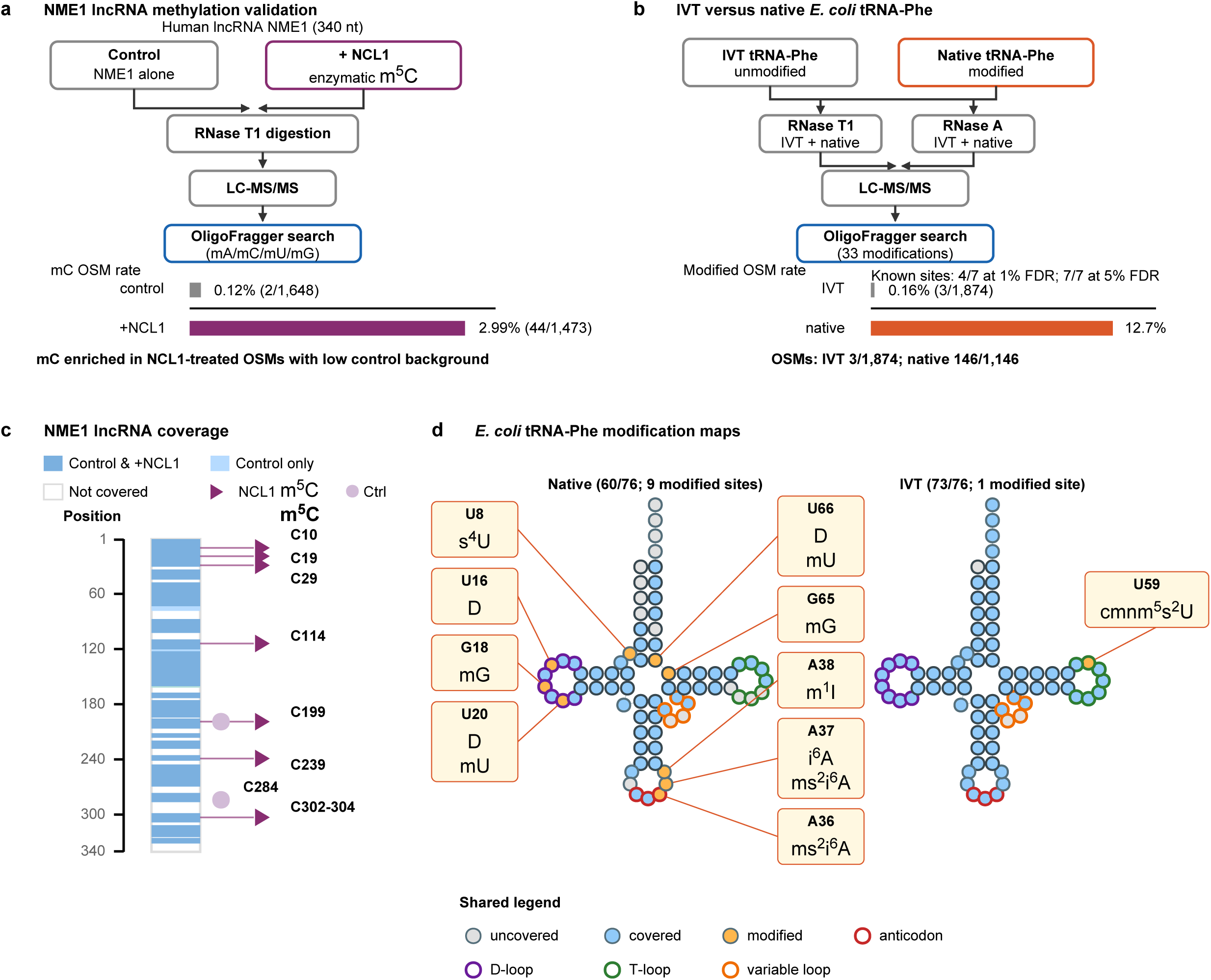
Validation of OligoFragger with controlled samples. **(a)** Reanalysis of the NASE NME1 lncRNA methylation-validation dataset and mC OSM rate for untreated control (1,648 OSMs, 2 with mC, 0.12%) and NCL1-treated samples (1,473 OSMs, 44 with mC, 2.99%). NCL1 treatment produced a strong enrichment of methylcytidine-compatible assignments in a focused methylated-base search. **(b)** IVT versus native *E. coli* tRNA-Phe-GAA validation workflow and modified-OSM rate; IVT has 3 non-suspicious modified OSMs among 1,874 total OSMs (0.16%) at 1% FDR, whereas native tRNA has 146 among 1,146 OSMs (12.7%). The tRNA-Phe validation was searched with a broad modification panel rather than a list restricted to the expected tRNA-Phe modifications. **(c)** Compact NME1 lncRNA sequence coverage map: each nucleotide of the 340-nt NME1 sequence is shown with background colored by detection in control and/or NCL1-treated conditions; detected mC sites are marked as filled purple triangles for NCL1-treated samples and lavender circles for control. **(d)** IVT and native *E. coli* tRNA-Phe-GAA cloverleaf diagrams showing nucleotide-level MS/MS coverage and detected modification sites; native tRNA-Phe recovered 4 of 7 annotated MS-detectable sites at 1% FDR and all 7 at 5% FDR, whereas pseudouridine at position 55 is mass-silent and excluded. Suspicious modification assignments are excluded throughout.

In the NCL1-treated runs, OligoFragger identified 1,473 OSMs at 1% FDR, of which 44 carried non-suspicious modifications. All 44 were assigned mC (the broad methylcytidine category covering m^5^C and related methylations), consistent with the known substrate specificity of NCL1 (Fig. 2a, c). The mC OSM rate increased from 0.12% in the untreated control to 2.99% after NCL1 treatment, showing that OligoFragger can recover the expected methylcytidine signals while suppressing competing methylated-base assignments.

#### IVT vs. native E. coli tRNA-Phe-GAA

As a more stringent validation, we compared modification profiles between in vitro transcribed (IVT) and native *E. coli* tRNA-Phe-GAA (anticodon GAA). IVT tRNA is synthesized without the cellular modification machinery and is devoid of post-transcriptional modifications, serving as a negative control. Both IVT and native tRNA were subjected to complete digestion with RNase T1 and RNase A separately, generating four runs (Fig. 2b).

*E. coli* tRNA-Phe-GAA carries 7 annotated modification sites detectable by mass spectrometry (s^4^U at position 8; D at positions 16 and 20; ms^2^i^6^A at position 37; m^7^G at position 46; acp^3^U at position 47; m^5^U at position 54), plus pseudouridine at position 55 which is mass-silent and not detectable by MS. The IVT and native tRNA-Phe datasets were searched with a broad modification panel of 33 candidate RNA modifications (Supplementary Table 2), rather than a list restricted to the known tRNA-Phe modifications. At 1% FDR, after excluding suspicious modification assignments, OligoFragger identified 4 of the 7 detectable annotated sites in native tRNA-Phe: s^4^U8, D16, D20 and ms^2^i^6^A37 (Fig. 2d). At 5% FDR, all 7 detectable annotated sites were recovered, including mG46 and acp^3^U47 from the short trinucleotide [mG][acp^3^U]C and mU54 from the T-loop.

Critically, IVT tRNA is synthesized without cellular modification machinery and does not contain post-transcriptional modifications by design. After excluding modification sites flagged as suspicious, only 3 of 1,874 IVT tRNA-Phe OSMs at 1% FDR retained a modification assignment (0.16%), compared with 146 of 1,146 OSMs in native tRNA (12.7%) (Fig. 2b). Thus, even when a large modification panel was searched, OligoFragger preserved a near-zero modification rate in the IVT negative control while recovering annotated modification sites in native tRNA.

### Comprehensive *E. coli* tRNA modification profiling

*E. coli* tRNA is among the most thoroughly characterized epitranscriptomes, with extensive site-level reference annotations available from a tRNAdb-derived *E. coli* modification dataset compiled by Kimura et al. ^33^, making it an ideal benchmark for comprehensive modification profiling. We analyzed *E. coli* total tRNA using a structure-protected RNase digestion strategy adapted from our previous total *Saccharomyces cerevisiae* tRNA LC-MS/MS workflow^38^. In that approach, folded tRNA structure restricts RNase cleavage to primarily single-stranded regions of the tRNA and increases the length, and thus the uniqueness of oligonucleotide products from total-tRNA mixtures. Here, total *E. coli* tRNA was digested with either RNase T1 or RNase A to generate overlapping oligonucleotide fragments with up to 10 missed cleavages, maximizing sequence coverage and enabling modification localization through corroborating evidence from overlapping sequences.

We first asked whether partial digestion could provide enough sequence information for repertoire-scale tRNA profiling while remaining searchable by OligoFragger. At 1% FDR, OligoFragger identified 4,539 OSMs from the combined RNase T1 (2,506) and RNase A (2,033) datasets. These 4,539 OSMs corresponded to 883 unique sequences — where two OSMs mapping to the same genomic locus but carrying different modifications count as distinct sequences. Of these, 766 (87%) could be assigned to a single tRNA anticodon species without ambiguity; the remaining 13% mapped to sequence regions shared across multiple isoacceptors. The accepted oligonucleotides ranged from 5 to 25 nucleotides, with a median length of 12 nt (Fig. 3a), and included both fully cleaved and missed-cleavage-containing products from the RNase T1 and RNase A datasets (Fig. 3b). Mapping these fragments onto Sprinzl positions produced broad coverage across all 39 *E. coli* tRNA isoacceptors (Fig. 3c; Supplementary Fig. 1): median isoacceptor coverage was 78.8%, representing, to our knowledge, the most extensive sequence coverage of the *E. coli* tRNA repertoire reported by oligonucleotide LC–MS/MS to date. Ile-GAU, Pro-UGG and Thr-GGU reached 100% coverage, and Sec-UCA was the lowest-covered isoacceptor (17.5%). Thus, the combined partial-digestion strategy generated long, overlapping sequence fragments across the *E. coli* tRNA repertoire and OligoFragger successfully searched these products, including those with missed cleavages, to support high-coverage modification profiling.

**Figure 3.**
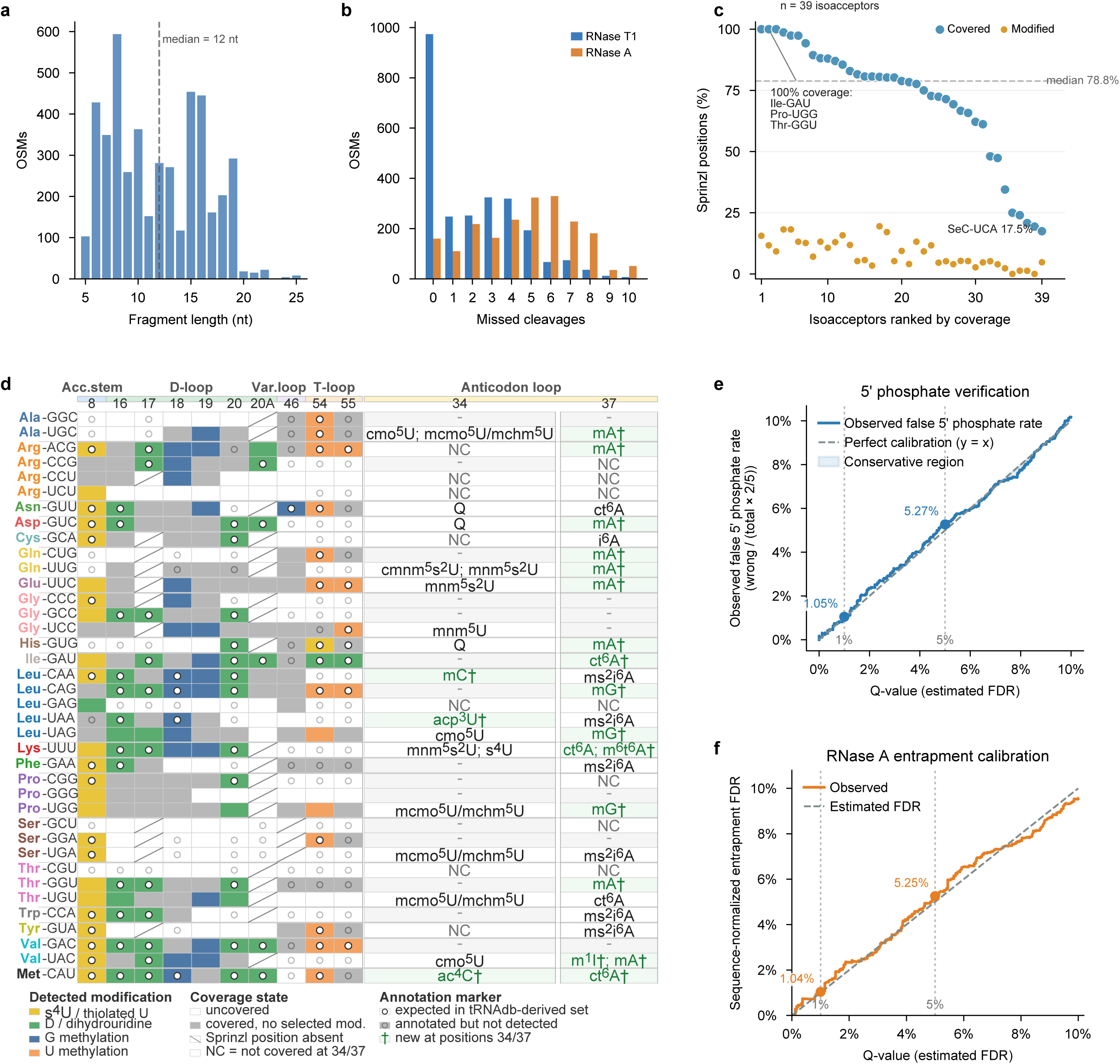
Comprehensive modification profiling of *E. coli* tRNA. **(a)** Fragment length distribution of accepted OSMs from the combined RNase T1 and RNase A searches (4,539 total at 1% FDR); dashed line indicates the median length of 12 nt. **(b)** Missed-cleavage distribution for RNase T1 (blue) and RNase A (orange) identifications shown as side-by-side bars. **(c)** Ranked Sprinzl-position coverage across 39 *E. coli* tRNA isoacceptors. Blue points show the percentage of covered Sprinzl positions, orange points show the percentage carrying a detected non-suspicious modification, and the dashed line marks the median coverage of 78.8%. **(d)** Integrated selected-position modification map. The left matrix summarizes selected structural Sprinzl positions; the right table shows exact anticodon-loop positions 34 and 37. Cell color indicates detected modification class, gray indicates coverage without a selected modification, open circles mark expected tRNAdb-derived annotations, gray circles mark annotated but undetected sites, and green daggers mark candidate sites not present in the reference annotation set at positions 34 or 37. **(e)** Independent 5’ phosphate verification showing the observed false 5’ phosphate rate as a function of nominal q-value; the observed rates were 1.05% and 5.27% at the 1% and 5% thresholds. **(f)** Entrapment FDR calibration from an RNase A combined *E. coli*/*S. cerevisiae* tRNA database search; yeast-exclusive matches provide empirical false positives after sequence-space normalization, yielding 1.04% and 5.25% estimates at the 1% and 5% thresholds.

To validate the accuracy of the modification assignments, we next examined whether the detected sites followed known positional patterns in *E. coli* tRNA. We examined two patterns: recurrent modifications that appear at conserved structural positions across many tRNA types, and highly isoacceptor-specific modifications concentrated at functionally specialized positions such as Sprinzl 34 and 37. In the selected-position map (Fig. 3d), modification calls were not randomly distributed. s^4^U was enriched at position 8, dihydrouridine and related calls appeared in the D-loop, methylated guanosines were concentrated in the D-loop and variable-loop/T-arm region, and U methylation was repeatedly detected at position 54. This recovery of known structural and decoding hotspots provides a repertoire-level accuracy check before considering sites beyond the current annotation set.

The anticodon loop provided a more stringent test of whether OligoFragger could resolve isoacceptor-specific modifications. Positions 34 and 37 are among the most functionally consequential tRNA modification sites, and the integrated Fig. 3d table showed that most detected calls at these positions were also annotated in the tRNAdb-derived reference set. At position 34, these included queuosine in Asn-GUU, Asp-GUC and His-GUG; cmo^5^U-family chemistry in Ala-UGC, Leu-UAG and Val-UAC; mcmo^5^U/mchm^5^U-family chemistry in Ala-UGC, Pro-UGG, Ser-UGA and Thr-UGU; mnm^5^s^2^U/m^5^s^2^U-family chemistry in Gln-UUG, Glu-UUC and Lys-UUU; mnm^5^U in Gly-UCC; and s^4^U in Lys-UUU. At position 37, database-annotated matches included i^6^A in Cys-GCA, ms^2^i^6^A in Leu-CAA, Leu-UAA, Phe-GAA, Ser-UGA, Trp-CCA and Tyr-GUA, and ct^6^A in Asn-GUU, Lys-UUU and Thr-UGU. This agreement supports the site-level accuracy of OligoFragger in a dense *E. coli* tRNA search space.

After establishing concordance with known modification patterns, we used the same analysis to identify candidate sites absent from the tRNAdb-derived reference annotation set. The strongest recurring example was mA-family evidence at Sprinzl position 37 in nine isoacceptors (Ala-UGC, Arg-ACG, Asp-GUC, Gln-CUG, Gln-UUG, Glu-UUC, His-GUG, Thr-GGU and Val-UAC, where the Val-UAC call was unresolved as m^1^I/mA). Other position-37 candidates marked as new in Fig. 3d included mG in Leu-CAG, Leu-UAG and Pro-UGG, ct^6^A in Ile-GAU and Met-CAU, and m^6^t^6^A in Lys-UUU. Figure 3d also highlights three position-34 candidates: mC in Leu-CAA, acp^3^U in Leu-UAA and ac^4^C in Met-CAU. To show the spectrum-level evidence behind these candidates, we selected two annotated examples: a position-37 anticodon-loop fragment, CCCU[mnm^5^s^2^U]UC[mA]CG, in which the candidate mA-family signal is observed alongside mnm^5^s^2^U at the wobble position (Supplementary Fig. 2), and a longer RNase A product, AAGGGAUU[acp^3^U]AA[ms^2^i^6^A]AUCCCU, supporting the candidate acp^3^U signal in Leu-UAA at position 34 (Supplementary Fig. 3). Repertoire-level summaries in Supplementary Figs. 4 and 5 show how these candidates sit within the broader *E. coli* modification landscape, with the full Sprinzl-position matrix and searched modification panels provided in Supplementary Tables 1 and 2. The candidate placements at decoding-adjacent positions make them useful examples of OligoFragger’s discovery potential, but these calls remain as spectrum-supported hypotheses that require orthogonal MS/MS or biochemical confirmation.

Finally, we tested whether the expanded modified-tRNA search space remained statistically controlled after accepting these high-coverage identifications. We used two independent calibration strategies. First, we exploited a fundamental property of enzymatic RNA digestion: in an intact tRNA molecule, a 5’ phosphate group exists only at the first nucleotide, whereas internal RNase cleavage products carry 5’-hydroxyl termini. Thus, a 5’-phosphate assignment on an internal fragment provides an empirical mass-coincidence false-positive readout that is independent of the target-decoy mechanism. Among all 6,955 non-decoy OSMs in the combined RNase T1 + RNase A dataset, 1,466 (21.1%) carried a 5’ phosphate assignment. After accounting for the two 5’-phosphate-containing precursor mass states among the five mass-offset states evaluated by OligoFragger, the observed false 5’ phosphate rate closely tracked the nominal q-value across the full range (Fig. 3e): 1.05% at the 1% FDR threshold and 5.27% at the 5% threshold. Second, we searched the RNase A data against a combined *E. coli*/*Saccharomyces cerevisiae* tRNA database and used yeast-exclusive matches as entrapment false positives. Assignments shared between *E. coli* and yeast parent sequences were counted as *E. coli*-compatible target hits rather than entrapment hits, and the entrapment rate was normalized by the ratio of the *E. coli*-only to yeast-only digested sequence spaces (11,967/10,360; Methods). The sequence-normalized entrapment estimates were 1.04% at the 1% FDR threshold and 5.25% at the 5% threshold (Fig. 3f). Together, the 5’ phosphate and entrapment analyses show that OligoFragger’s false discovery rate remains well controlled during comprehensive modified-tRNA profiling.

### Human tRNA benchmarking and modification profiling

#### Independent benchmark against NASE

We next used the human HAP1 tRNA dataset originally analyzed with the NucleicAcidSearchEngine (NASE)^31^ as an external benchmark for OligoFragger. This dataset presents several analytical challenges: RNase T1 digestion was close to complete (91.6% of OSMs with zero missed cleavages; Fig. 4a, inset), producing short fragments (median, 10 nt; Fig. 4a). The data were also acquired using higher-energy collisional dissociation, which can generate additional RNA fragmentation, including internal and low-mass ions^39^. Because the >200 human tRNA genes and isoacceptors are highly similar, these short, modified oligonucleotides often share composition and precursor mass with multiple plausible targets. The result is a difficult search space in which many target and decoy candidates, including isobaric or near-isobaric modification variants, compete for spectra with incomplete or noisy fragment evidence. OligoFragger’s spectrum-first strategy is designed for this setting because graph-derived routes and diagnostic ions constrain candidate modifications before full scoring.

**Figure 4.**
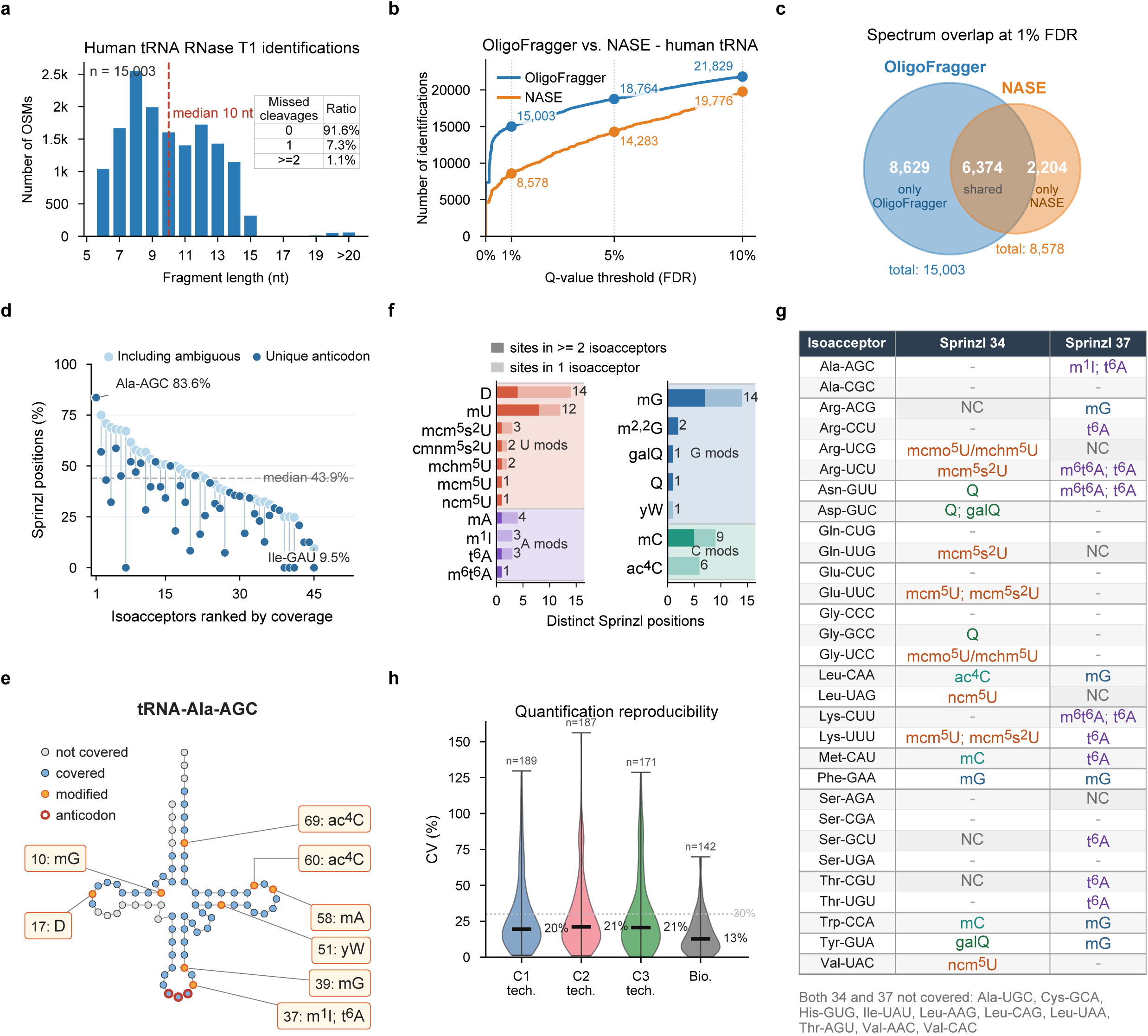
Human tRNA profiling with OligoFragger and comparison with NASE. **(a)** Identification statistics for the human HAP1 tRNA RNase T1 dataset (9 replicates, 1% FDR; n=15,003 target OSMs). Main plot: fragment length distribution, with dashed line indicating the median of 10 nt. Inset: missed-cleavage ratios, showing that 91.6% of OSMs have zero missed cleavages. **(b)** Cumulative target OSM counts plotted against q-value threshold for OligoFragger (blue) and NASE (orange). OligoFragger identifies 15,003 vs. 8,578 OSMs at 1% FDR (1.75-fold) and 21,829 vs. 19,776 OSMs at 10% FDR (1.10-fold). **(c)** Proportional Venn diagram of spectra identified at 1% FDR: 6,374 shared (matched by source file + scan number), 8,629 OligoFragger-exclusive and 2,204 NASE-exclusive. **(d)** Per-isoacceptor Sprinzl-position coverage by accepted OligoFragger sequences, with paired dots showing coverage calculated with ambiguous anticodon-mapped OSMs included or restricted to unique-anticodon OSMs. **(e)** Cloverleaf view of tRNA-Ala-AGC, the most-covered human isoacceptor. Circles represent Sprinzl positions: gray, no accepted fragment coverage; blue, covered without a detected modification; orange with dark outline, non-suspicious detected modification; red outline, anticodon positions 34-36. Boxed labels show the modified Sprinzl position and modification call. **(f)** Overall human tRNA modification-type landscape. Bars show the number of distinct Sprinzl positions detected for each modification type, grouped by parent nucleobase and split by whether a site was supported by one or at least two isoacceptors. **(g)** Anticodon-loop modification and coverage table for exact Sprinzl positions 34 and 37. Cells list non-suspicious OligoFragger modification calls, "-" indicates sequence coverage without a detected modification and "NC" indicates no accepted fragment covered the Sprinzl position for that isoacceptor. Semicolons separate multiple calls at the same position; the slash in mcmo^5^U/mchm^5^U denotes unresolved isobaric modification evidence. **(h)** OSM-level quantification reproducibility across 9 runs (3 biological clones x 3 technical replicates). Violin plots show the distribution of coefficients of variation (CV) computed from median-normalized linear intensities for sequences detected in all replicates within each group. Technical CVs have median values of 19.5%, 21.1% and 20.6% for clones 1-3, respectively; the biological CV across clone means has a median of 12.7%. Dashed line: 30% CV reference.

Under this framework, OligoFragger identified 15,003 target OSMs at 1% FDR, representing 408 unique modified or unmodified sequences and 190 unique anticodon-defined fragment loci (Fig. 4a). Of these sequences, 288 (70.6%) mapped uniquely to a single anticodon-defined tRNA species and 120 mapped to regions shared by multiple isoacceptors, a lower unique-mapping rate than observed for the longer partial-digestion *E. coli* fragments. When accepted OSM counts were plotted against the q-value threshold, OligoFragger recovered more target OSMs than NASE under stringent FDR control (Fig. 4b): 15,003 versus 8,578 OSMs at 1% FDR (1.75-fold). The advantage narrowed at a more permissive 10% FDR threshold (21,829 versus 19,776 OSMs; 1.10-fold), consistent with the interpretation that spectrum-guided candidate pruning is most beneficial when decoy competition imposes strict score cutoffs. This gain was achieved despite OligoFragger not including the sodium and potassium adduct search channels used by NASE.

We examined the overlap between tools to assess agreement. Matching spectra by source file and scan number at 1% FDR, 6,374 spectra were identified by both tools, whereas 8,629 were identified only by OligoFragger and 2,204 only by NASE (Fig. 4c). Among shared spectra with sequence-concordant assignments, modification calls largely agreed after nomenclature normalization, indicating that OligoFragger’s increased recovery did not arise from globally reinterpreting the spectra that both tools can already assign. Instead, the principal gain was recovery of additional spectra in a highly modified, short-fragment search space where many plausible but incorrect candidates otherwise compete for the top score.

#### Human tRNA modification landscape

We first examined how broadly the accepted OligoFragger identifications covered the human tRNA repertoire after mapping fragments onto universal Sprinzl positions. Coverage was uneven across isoacceptors (Fig. 4d). The highest coverage was observed for tRNA-Ala-AGC, with 83.6% of Sprinzl positions covered by sequences that mapped uniquely to one anticodon; this isoacceptor is shown at nucleotide resolution in Fig. 4e. Across detected isoacceptors, the median unique-anticodon coverage was 43.9%. Including ambiguous anticodon-mapped fragments increased apparent coverage for many tRNAs, but coverage dropped substantially when the analysis was restricted to uniquely mapped fragments. This drop is expected for human tRNA data because many human tRNA isoacceptors and isodecoders are highly similar, and the short RNase T1 fragments in this dataset often cannot distinguish among closely related tRNA sequences.

The human search was constrained to a curated human tRNA modification list containing 33 candidate modification types. After suspicious assignments were removed, OligoFragger detected 20 of these 33 candidate types at 50 Sprinzl positions across 43 human tRNA isoacceptors (Fig. 4f). The remaining searched modification types were absent from the accepted non-suspicious target set and are not interpreted further in the main text. To illustrate the spectrum-level evidence behind these calls, Supplementary Fig. 6 shows a t^6^A-containing fragment, ACUCUU[t^6^A]AUCUCAG, in which the t^6^A diagnostic ion accompanies backbone-fragment evidence across the oligonucleotide, while Supplementary Fig. 7 shows that internal non-labile mU residues in AU[mU]CCCAU[mU]CUUG can still generate high intensity modified single-nucleotide ions. The underlying isoacceptor-level calls and coverage states are reported in Supplementary Table 3. We interpreted these calls against prior knowledge of human tRNA modification biology, especially the enrichment of functional modifications in the anticodon loop and the established roles of positions 34 and 37 in decoding and reading-frame maintenance^3,40,41^.

The Sprinzl-position summaries showed that the detected landscape followed the modular architecture of mature tRNAs rather than a random distribution of mass shifts (Supplementary Figs. 8 and 9). Structural modifications were distributed through the acceptor stem, D arm, variable loop and T arm, whereas modifications associated with decoding were concentrated in the anticodon loop (Fig. 4g). In the modification-type summary, methylated residues were the most widely distributed class: mA, mU, mC, mG and m^2,2^G collectively appeared at 38 Sprinzl positions across 40 isoacceptors (Fig. 4f). The strongest single-position signals included canonical T-loop and neighboring structural sites such as mA58, mU54 and mU55, together with recurrent mG7, mG37 and mC49 (Supplementary Figs. 8 and 9). These sites provide an internal positive control that the workflow recovers the expected conserved modification framework before asking what is specific to individual anticodons^3,41^.

The clearest specificity emerged at Sprinzl positions 34 and 37, the two anticodon-loop positions that directly tune decoding (Fig. 4g). Position 37 modifications are expected to stabilize the anticodon loop and reduce frameshifting, but their distribution is still isoacceptor-selective^3,40,41^. After applying the position-37 rule described in Methods, labile t^6^A-family signals were assigned to 11 isoacceptors: Ala-AGC, Arg-CCU, Arg-UCU, Asn-GUU, Lys-CUU, Lys-UUU, Met-CAU, Ser-GCU, Thr-AGU, Thr-CGU and Thr-UGU. This set is biologically coherent because t^6^A37 is a conserved feature of tRNAs decoding ANN codons^40^. The methylated t^6^A form m^6^t^6^A was assigned to Arg-UCU, Asn-GUU and Lys-CUU, suggesting further modification of t^6^A in a subset of isoacceptors.

A second position-37 pattern in Fig. 4g was mG. OligoFragger detected mG at Sprinzl 37 in several isoacceptors, including Arg-ACG, Arg-CCG, Arg-UCG, Leu-CAA, Phe-GAA, Trp-CCA and Tyr-GUA, consistent with the broader recurrence of position-37 methylation in the human tRNA landscape. OligoFragger also reported an i^6^A-containing Ser-family assignment corresponding to the ambiguous Ser-AGA/Ser-CGA/Ser-UGA fragment group, consistent with prior evidence that human TRIT1 acts on a limited subset of anticodon-loop substrates^42^. Because this fragment is shared among closely related Ser isoacceptors, we do not assign the i^6^A signal to a single anticodon.

At the wobble position, the biology was different. Position 34 modifications directly shape codon recognition, especially for U34- and G34-containing anticodons^3,40,41^. OligoFragger recovered expected U34 chemistry in several split-box decoding tRNAs: mcm^5^s^2^U at Sprinzl 34 in Arg-UCU, Gln-UUG, Glu-UUC and Lys-UUU, and mcm^5^U at the same position in Glu-UUC and Lys-UUU. These 5-substituted and thiolated U34 derivatives are known to tune wobble decoding and translation efficiency^40^. Q-family chemistry provided a second expected wobble class: Q was detected at Sprinzl 34 in Asn-GUU and Asp-GUC, whereas galQ was detected at Sprinzl 34 in Asp-GUC and Tyr-GUA, matching the canonical role of queuosine-family modifications in NAY-decoding tRNAs^40,43^. Additional Sprinzl-34 signals, including Q in Gly-GCC, mchm^5^U, ncm^5^U, mC, ac^4^C and mG, represent candidate sites where OligoFragger may help refine isoacceptor-specific modification maps.

The dataset also contained signals outside the canonical anticodon-loop positions. Some were expected structural modifications, such as recurrent m^2,2^G at positions known to participate in tRNA core architecture, reinforcing the idea that OligoFragger captures both decoding and folding-associated chemistry. Other signals were not clearly represented in current human tRNA annotations but were repeatedly observed in defined sequence contexts. The most notable example was an Arg-associated Q signal at Sprinzl 49: nine OSMs contained the fragment AUU[Q]UG, expanded to Arg-CCG and Arg-CCU, with q-values from 0.0052 to 0.0099. We also observed ac^4^C at atypical sites beyond the commonly discussed position-34 and D-arm contexts, including Thr-AGU at Sprinzl positions 25 and 27 and Ala-AGC at Sprinzl positions 60 and 69; these sites are potentially interesting and warrant follow-up.

#### Label-free quantification reproducibility

Finally, because the public human tRNA dataset does not contain a perturbation condition, we treated quantification here as a reproducibility test rather than as evidence for biological regulation. Integration with IonQuant allows us to measure the chromatographic MS1 signal associated with each identified oligonucleotide, providing an intensity-based estimate of its relative abundance. These measurements allow the abundance of the same modified or unmodified oligonucleotide form to be compared across samples. We evaluated the reproducibility of these measurements across nine replicate runs (Fig. 4h; Supplementary Fig. 10). For each unique sequence per run, the maximum quantified intensity was retained, yielding a sequence by run intensity matrix of 382 unique quantified sequences (306 modified, 76 unmodified). After median normalization, the coefficient of variation (CV) was computed for sequences detected in all three technical replicates within each biological clone and, separately, for sequences detected in all nine runs (n=142). Technical CVs had median values of 19.5%, 21.1% and 20.6% across the three clones, with approximately half of sequences below the 20% threshold - comparable to what is routinely observed in label-free proteomics experiments on similar instruments. The biological CV, computed from clone means, was lower (median 12.7%), indicating that clone-to-clone variation in HAP1 cells contributes less noise than the technical variability of the LC-MS/MS measurement itself. Together, these results show that OligoFragger identifications can support reproducible MS1-based quantification across biological and technical replicates; differential biological claims would require matched experimental perturbations.

## DISCUSSION

For decades, the identification of highly modified RNAs by oligonucleotide MS/MS has been limited not only by spectral complexity but also by the computational cost of considering many possible modifications in one search. This problem is especially acute for tRNAs, where a realistic search may require dozens of candidate modified residues and multiple modifications within a single oligonucleotide. In sequence-enumeration workflows, these possibilities behave like many variable modifications, expanding both target and decoy candidate spaces and reducing sensitivity under FDR control. OligoFragger addresses this challenge by reversing the order in which modified candidates are considered. Instead of enumerating modified sequence variants before spectrum matching, it first uses the experimental spectrum to build candidate sequence routes and introduces modification hypotheses only when fragment evidence supports them. This spectrum-first design keeps the database in unmodified sequence space, supports target-decoy FDR estimation and allows broad modification panels to be searched while retaining sensitivity for densely modified RNA oligonucleotides.

The control experiments showed that this search strategy can detect true RNA modifications while maintaining low false-positive rates. In reanalysis of the NASE lncRNA methyltransferase dataset and in IVT/native tRNA-Phe experiments, OligoFragger recovered expected modification signals in modified samples while producing near-zero non-suspicious modification assignments in negative controls. In comprehensive *E. coli* tRNA profiling, OligoFragger achieved broad isoacceptor coverage, matched many modification patterns in the tRNAdb-derived reference annotations, and maintained calibrated error control in a dense modified-tRNA search space, as supported by independent 5’ phosphate and entrapment analyses.

Application to the independent human HAP1 tRNA dataset further showed that OligoFragger improves recovery under stringent FDR control while maintaining agreement with an established search engine. Relative to NASE, OligoFragger identified 1.75-fold more target OSMs at 1% FDR, and spectra assigned the same sequence by both tools showed strong agreement in modification calls after nomenclature normalization. The accepted identifications allowed us to map fragments to Sprinzl positions across the tRNA repertoire, examine position-specific modification patterns, identify candidate sites beyond current annotation sets, and perform reproducible label-free quantification across biological and technical replicates.

Several limitations should guide interpretation of OligoFragger results. First, some RNA modifications remain difficult to distinguish by mass spectrometry alone. Mass-silent modifications such as pseudouridine cannot be separated from unmodified uridine by mass shift alone in standard oligonucleotide MS/MS. Second, identification depth and localization confidence still depend on the experimental MS workflow, including digestion strategy, chromatographic separation, fragmentation efficiency and acquisition depth. Together, these results establish OligoFragger as a practical and statistically controlled platform for large-scale RNA oligonucleotide MS/MS search, modification mapping, and quantitative profiling. OligoFragger provides command-line and graphical user interfaces, support for multiple MS data formats, and customizable modification databases, supporting broader use of mass spectrometry for RNA modification analysis.

## METHODS

### In silico digestion and decoy generation

Input RNA sequences are digested in silico according to the user-selected cleavage rule, with configurable missed cleavages and length filtering. Shared digestion products are deduplicated while retaining parent identifiers. The resulting database remains in unmodified sequence space; searched modifications and terminal states are introduced later during spectrum-graph matching and precursor validation.

For FDR estimation, decoy sequences are generated from the target sequence space by shuffling, dinucleotide shuffling or reversal. Decoys preserve the selected sequence-composition constraint, are checked to avoid collisions with target and existing decoy entries, and are marked with a ‘DECOY_’ prefix.

### Overview of the search algorithm

The central algorithmic innovation in OligoFragger is a spectrum graph-guided database search. Rather than scoring every database candidate against every spectrum (which scales poorly with large modification databases), OligoFragger first extracts compositional information directly from each experimental spectrum and uses it to restrict the candidate space. The search proceeds through the following stages for each MS/MS spectrum:

**Stage 1 — Deisotoping**. Raw MS/MS peaks are deisotoped and decharged to generate a compact list of neutral-mass peaks for graph construction. Each deisotoped peak retains its representative mass, charge state and intensity.

**Stage 2 — Spectrum graph construction**. Deisotoped peaks are converted into a directed spectrum graph. Edges connect peak pairs whose mass differences match canonical or enabled modified residues, so routes through the graph represent sequence and modification evidence supported by the observed fragments.

**Stage 3 — Node labeling**. Graph nodes are matched to a theoretical library of low-mass one- and two-nucleotide RNA fragment ions, including enabled modified residues. Including two-nucleotide ions allows routes to begin from early fragment evidence even when the first single-nucleotide ion is missing, which is common for longer oligonucleotides. This step links spectrum-graph routes to interpretable backbone-ion evidence; the ion-label convention is defined in Supplementary Note 1.

**Stage 4 — Diagnostic ion filtering (optional).** When enabled, diagnostic ion filtering is applied before spectrum-graph construction. OligoFragger first scans the raw MS/MS peaks for modification-specific low-mass evidence: fixed diagnostic m/z values are used for labile modifications such as t^6^A-family residues, whereas other modifications are tested using theoretical one-nucleotide fragment ions generated for the modified residue. Only modifications with diagnostic support are retained as possible graph-edge labels and node-label ions for that spectrum. Labile diagnostic-ion modifications are also tracked separately, because their diagnostic base-loss signal may support a precursor-mass-compatible assignment even when the modified residue is not retained in conventional backbone fragment ions.

**Stage 5 — Route extraction.** Starting from graph nodes that match theoretical one- or two-nucleotide ions, OligoFragger follows connected residue edges to assemble spectrum-supported sequence routes. Routes are retained up to a fixed maximum length or until no further compatible edge is available, and each route records its starting ion evidence, inferred sequence composition and matched-fragment support. These routes are used to filter candidate database sequences and are later evaluated as prefix or suffix evidence during modification assignment and scoring.

**Stage 6 — Candidate matching and precursor validation.** Database sequences are first filtered by an unmodified-mass window around the precursor mass. Spectrum-supported routes are then tested as prefix and suffix evidence against each compatible unmodified sequence, and the matched routes are used to generate focused modification-aware assignments. Each assignment is retained only if its theoretical mass satisfies precursor validation after isotope-error correction and terminal-state offsets are considered. The terminal offsets include no offset, HPO3, HPO3-H2O, 2HPO3 and 2HPO3-H2O, representing alternative phosphate and water-loss states evaluated during validation rather than pre-expanded as separate digestion products. When terminal-ion evidence is informative, the allowed offset state is restricted according to the ion-label convention and terminal-state model described in Supplementary Note 1.

**Stage 7 — Scoring.** Surviving candidates are scored using the matched graph nodes from the selected prefix and suffix routes. For each candidate, OligoFragger sums the mass-accuracy-weighted intensities of nonredundant matched route nodes and normalizes this evidence against high-intensity local peaks in the decharged spectrum. The final OligoFragger score rewards both the number of unique matched route nodes and the relative intensity support for those nodes:

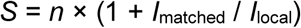

where *n* is the number of nonredundant matched graph nodes, *I*_matched_ is the summed weighted intensity of matched route evidence, and *I*_local_ is the summed intensity of selected local high-intensity peaks around the matched node masses. Higher scores therefore require both broader fragment-route coverage and stronger spectrum-level intensity support.

**Stage 8 — Ranking.** Candidates are sorted by descending score, and the top *k* matches per spectrum are retained (default *k* = 1). Proper ranking is assigned accounting for ties.

### False discovery rate estimation

FDR is estimated using target-decoy competition. For each spectrum, the best-scoring target or decoy match is retained, and retained matches are sorted by score. At each score threshold, the FDR is calculated as:

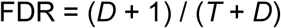

where *D* and *T* are the cumulative numbers of decoy and target hits at or above the threshold. q-values are assigned as the minimum FDR observed at each score or worse, and FDR is calculated independently for each input file before filtered results are combined.

### Suspicious modification annotation

OligoFragger implements a spectrum-based suspicious modification annotation system to flag potentially unreliable modification assignments. A modification assignment is flagged as suspicious when an alternative route with the same local sequence context but fewer modifications provides stronger local spectrum support, or when terminal modified dinucleotide evidence is better explained by a less-modified composition. This comparison is performed after final OSM selection: the less-modified route may better explain local fragment evidence, but it is not selected as the final assignment if the corresponding less-modified candidate cannot satisfy precursor-mass validation under the allowed isotope and terminal-state offsets. Thus, suspicious annotations identify weak site-level modification support within an otherwise retained precursor-compatible OSM. These flags are based on spectrum-derived support rather than external biochemical plausibility rules, and the site-level criteria are described in Supplementary Note 2. Suspicious flags do not remove the identification; downstream filtering decisions are left to the user.

### Label-free quantification

OligoFragger provides optional MS1-based label-free quantification through integration with the IonQuant library. For each identified OSM, the extracted ion chromatogram (XIC) is constructed from the raw MS1 data using the precursor m/z, retention time, and charge state. IonQuant performs peak detection and integration using configurable quality criteria; by default, at least 2 isotope peaks and 3 scans are required for a valid quantification. Each match is annotated with the resulting intensity value in the output. Quantified intensities across runs enable comparative analysis of modification stoichiometry between experimental conditions.

### Materials and sample preparation

#### Public NASE datasets

Two datasets from the NASE publication^31^ were reanalyzed with OligoFragger. The NME1 lncRNA dataset contains six LC-MS/MS runs: three from untreated NME1 lncRNA and three from NME1 lncRNA incubated with the NCL1 methyltransferase. The human tRNA dataset (PRIDE PXD016328) contains crude cellular tRNA isolated in three biological replicates from HAP1 tissue culture using an RNeasy kit; the short-RNA fraction, enriched for RNAs <200 nt and mostly composed of tRNA, was digested with RNase T1 and analyzed by nano-flow ion-pair LC-MS/MS in technical triplicate on a Q Exactive HF. In total, nine human tRNA LC-MS/MS runs were reanalyzed.

### IVT and native *E. coli* tRNA-Phe-GAA

The unmodified tRNA-Phe-GAA control was generated through run off in vitro transcription using a DNA oligonucleotide with the following sequence: mUmGG TGC CCG GAC TCG GAA TCG AAC CAA GGA CAC GGG GAT TTT CAA TCC CCT GCT CTA CCG ACT GAG CTA TCC GGG CCC CTA TAG TGA GTC GTA TTA. The transcription products were gel purified using 10% urea-PAGE. The IVT product excised from the gel after UV shadow visualization. IVT tRNA-Phe-GAA was gel extracted overnight at 4°C in 0.3 M NaOAc pH 5.2 and 2 mM EDTA pH 8.0 while shaking. The extraction buffer containing IVT tRNA-Phe-GAA was filtered to remove gel pieces and concentrated using a 10 kDa molecular weight cut off filter. The concentrated IVT tRNA Phe-GAA solution was ethanol precipitated overnight using 1/10^th^ volume 3 M NaOAc pH 5.2, 2 μL glycoblue, and at least 2 volumes of 100% ice cold ethanol. The IVT-tRNA-Phe-GAA was spun down at 10,000 xg for 30 minutes and the pellet was rinsed with 70% ice cold ethanol twice. The IVT-tRNA-Phe-GAA pellet was dried and reconstituted in RNase and DNase free water. tRNA-Phe-GAA was purified from *E. coli* HB101 containing pUC57-tRNA^phe^ provided by Yury Polikanov as previously described^44^.

Native and IVT tRNA-Phe-GAA were subjected to orthogonal complete digestions using RNase T1 (100 U/μg) and RNase A (100 ng/μg) as previously described^38^. Briefly, 9 μg of each tRNA was digested using RNase T1 or RNase A in 220 mM ammonium acetate (pH unadjusted) for 1 hour at 37°C. RNase digestion was quenched using phenol:chloroform:isoamyl alcohol. The digested RNA oligonucleotides were dephosphorylated in 1 hour at 37°C using 0.13 U/ug quick CIP (NEB). The digested oligonucleotides were ethanol precipitated and resuspended in 200 mM ammonium acetate (pH unadjusted).

### *E. coli total* tRNA RNase digestion

Total tRNA was extracted from wild-type *E. coli* K-12 as previously described^45^. Total-tRNA preparation and RNase digestion followed the structure-protected total-tRNA LC-MS/MS strategy previously developed for total *S. cerevisiae* tRNAs^38^, applied to *E. coli*. Briefly, total tRNA was subjected to RNase T1 (6 U/ug) or RNase A (6 ng/ug) with quick CIP (0.13 U/ug) digestion to generate overlapping oligonucleotide fragments for repertoire-scale mapping. The LC–MS/MS conditions have been described previously.

### Data analysis parameters

All searches used 20 ppm precursor and fragment mass tolerances, shuffled decoys, target-decoy FDR estimation and isotope-error correction from −3 to +1. Digested-sequence length filtering was generally 5 to 25 nucleotides; the purified tRNA-Phe validation searches used 3 to 25 nucleotides to retain short complete-digestion products. Missed-cleavage allowances were set according to the search workflow: up to 3 missed cleavages for the NME1 RNase T1 searches and purified tRNA-Phe RNase T1/RNase A searches, up to 5 for the human tRNA RNase T1 searches and up to 10 for the partial-digestion *E. coli* tRNA searches. Diagnostic-ion filtering was enabled for the NME1 and human tRNA searches and disabled for the purified tRNA-Phe, *E. coli* profiling and entrapment searches. During precursor validation, OligoFragger evaluated no offset, HPO3, HPO3-H2O, 2HPO3 and 2HPO3-H2O terminal mass states. Modification panels are listed in Supplementary Table 2; q-value thresholds used for reporting are stated in the corresponding Results sections. Unless otherwise stated, all modification-level analyses excluded sites marked as suspicious by OligoFragger.

### *E. coli* tRNA reference annotation comparison

For the site-level *E. coli* comparison in Fig. 3d, reference annotations were taken from the Supplementary Data 1 table reported by Kimura et al. ^33^, which contains reference *E. coli* tRNA sequences with modifications retrieved from tRNAdb and updated with literature-based sequence corrections or additions. We used this tRNAdb-derived table as the primary site-level annotation standard because it provides a directly usable curated reference set for the same organism and modification-survey context. MODOMICS was still used for modification definitions, residue masses, short names and biosynthetic or pathway grouping where indicated, but not as the primary site-level annotation source for the *E. coli* comparison.

### Terminal-state and entrapment calibration analyses

For the 5’ phosphate calibration in Fig. 3e, accepted non-decoy OSMs from the *E. coli* RNase T1 and RNase A profiling searches were sorted by q-value. A 5’ phosphate assignment was defined by the two terminal-offset states containing phosphate on both the 5’ and 3’ sides (2HPO3 and 2HPO3-H2O). Such assignments were treated as false terminal-state calls when the mapped fragment started downstream of the mature tRNA 5’ end, because internal RNase cleavage products are expected to carry 5’-hydroxyl termini. At each q-value threshold, the cumulative false 5’ phosphate count was divided by two-fifths of the cumulative non-decoy OSM count, reflecting the two 5’-phosphate-containing states among the five terminal-offset states evaluated by OligoFragger.

For the entrapment calibration in Fig. 3f, the RNase A *E. coli* spectrum file was searched against the combined *E. coli*/*S. cerevisiae* tRNA database. The *E. coli*-only RNase A digestion space contained 11,967 sequences, whereas the combined database contained 22,327 sequences; the yeast-only entrapment space was therefore 10,360 sequences. At each q-value threshold, yeast-only non-decoy assignments were counted as entrapment hits, while assignments whose parent list included both *E. coli* and yeast sequences were counted as *E. coli*-compatible target hits because the observed spectrum could be explained by a target-database sequence. The raw entrapment rate was calculated as yeast-only entrapment hits divided by *E. coli*-compatible target hits and then multiplied by 11,967/10,360 to account for the different searchable sequence-space sizes.

### Human tRNA Sprinzl mapping and filtering

For human tRNA biological summaries, accepted OSMs were first filtered to the stated FDR threshold and modification assignments marked as suspicious by OligoFragger were excluded from the modification-landscape analysis. OligoFragger fragments were mapped to universal Sprinzl positions using the human Sprinzl-numbering table. Semicolon-delimited assignments within the same amino-acid family were expanded to their constituent isoacceptors; assignments spanning mixed amino-acid families were excluded from isoacceptor-level summaries. OligoFragger-exclusive identifications were not treated as novel biology solely because they were exclusive to OligoFragger. Biological summaries used non-suspicious assignments supported by target-decoy q-values, spectrum-graph route evidence, diagnostic-ion support where applicable, and mappable Sprinzl context.

Labile t^6^A-family calls were handled with an explicit positional rule. A t^6^A-family call was assigned to Sprinzl position 37 only when the raw OSM itself covered Sprinzl 37 and the reference base at that position was A; otherwise the OSM was retained at the spectrum level but not used as definitive position-37 evidence. A representative annotated t^6^A-containing spectrum is provided in Supplementary Fig. 6 to illustrate this evidence model.

## Data availability

The public NASE datasets reanalyzed in this study are available through the PRIDE repository: the NME1 lncRNA methyltransferase dataset under accession PXD016308 and the human HAP1 tRNA dataset under accession PXD016328. The *E. coli* total tRNA RNase T1 LC–MS/MS data reanalyzed in this study were originally reported by Simcox et al. ^45^ and have been deposited in the PRIDE repository under accession PXD068701. Raw LC–MS/MS data for the IVT and native *E. coli* tRNA-Phe-GAA experiments (RNase T1 and RNase A digests) and the RNase A digests of *E. coli* total tRNA will be deposited in PRIDE before publication.

## Code availability

OligoFragger will be made freely available to non-profit academic institutions under an academic license from the University of Michigan. Commercial licensing will be available separately through Fragmatics.

## Acknowledgements

This work was supported in part by the National Institutes of Health grants R01-GM-094231 and U24-CA271037 (to A.I.N), R01-HG013876 (to K.S.K.), and T32-GM132046 (to K.M.S.).

