## Supplementary Information for "OligoFragger: spectrum graph-guided analysis of modified RNA oligonucleotides by mass spectrometry"

| **Table of Contents** | S1 |
|---|---|
| **Summary of Supplementary Tables** | S2 |
| **Supplementary Figure 1.** Sequence coverage per *E. coli* tRNA isoacceptor | S3 |
| **Supplementary Figure 2.** Annotated spectrum for a candidate mA-family signal at Sprinzl position 37 in E. coli tRNA | S4 |
| **Supplementary Figure 3.** Annotated spectrum for a candidate acp3U signal at Sprinzl position 34 in *E. coli* tRNA | S5 |
| **Supplementary Figure 4.** *E. coli* tRNA modification prevalence across Sprinzl positions | S6 |
| **Supplementary Figure 5.** *E. coli* tRNA modification distribution across Sprinzl positions | S7 |
| **Supplementary Figure 6.** Representative annotated spectrum for a t6A-containing human tRNA fragment | S8 |
| **Supplementary Figure 7.** Representative annotated spectrum showing internal single-nucleotide evidence for a non-labile human tRNA modification | S9 |
| **Supplementary Figure 8.** Human HAP1 tRNA modification prevalence by Sprinzl region | S10 |
| **Supplementary Figure 9.** Human HAP1 tRNA modification distribution across Sprinzl positions | S11 |
| **Supplementary Figure 10.** OSM-level quantification completeness for human HAP1 tRNA | S12 |
| **Supplementary Note 1.** Terminal-state-aware precursor validation and ion-label convention | S13 |
| **Supplementary Note 2.** Suspicious modification annotation | S14 |

**Supplementary Tables**

**Supplementary Table 1. *E. coli* tRNA Sprinzl-position modification matrix.**

**Supplementary Table 2. Complete list of modifications searched in each analysis.**

**Supplementary Table 3. Human tRNA Sprinzl-position modification and coverage matrix.**

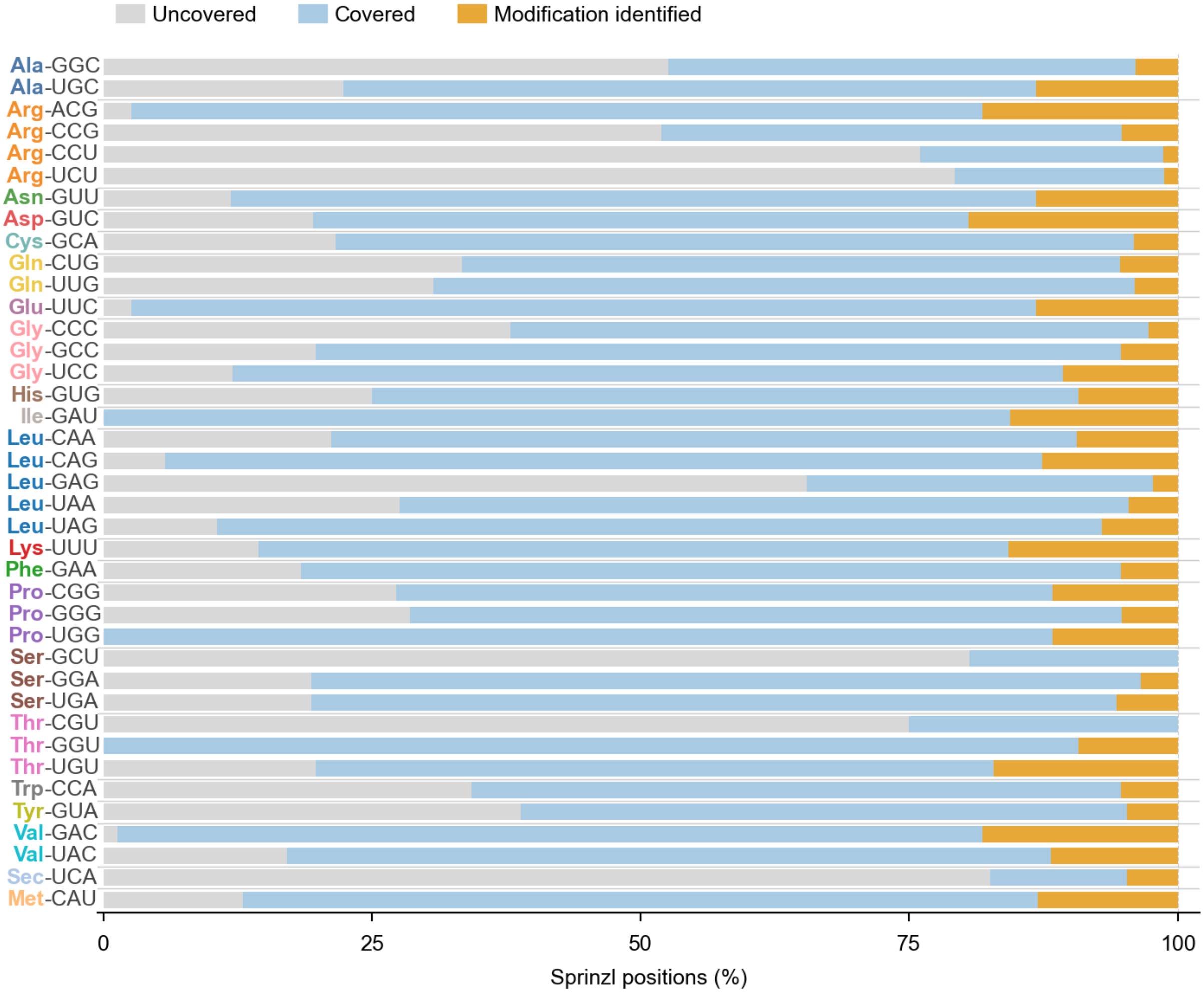


**Supplementary Figure 1. Sequence coverage per *E. coli* tRNA isoacceptor.**

Stacked horizontal bars show the fraction of Sprinzl positions th  at are uncovered, covered with no modification identified, or covered with at least one modification identified for each of the 39 tRNA isoacceptor species. Coverage ranges from 17.5% (Sec-UCA) to 100% (Ile-GAU, Pro-UGG and Thr-GGU). Identifications are from combined RNase T1 and RNase A searches at 1% FDR; suspicious modifications are excluded

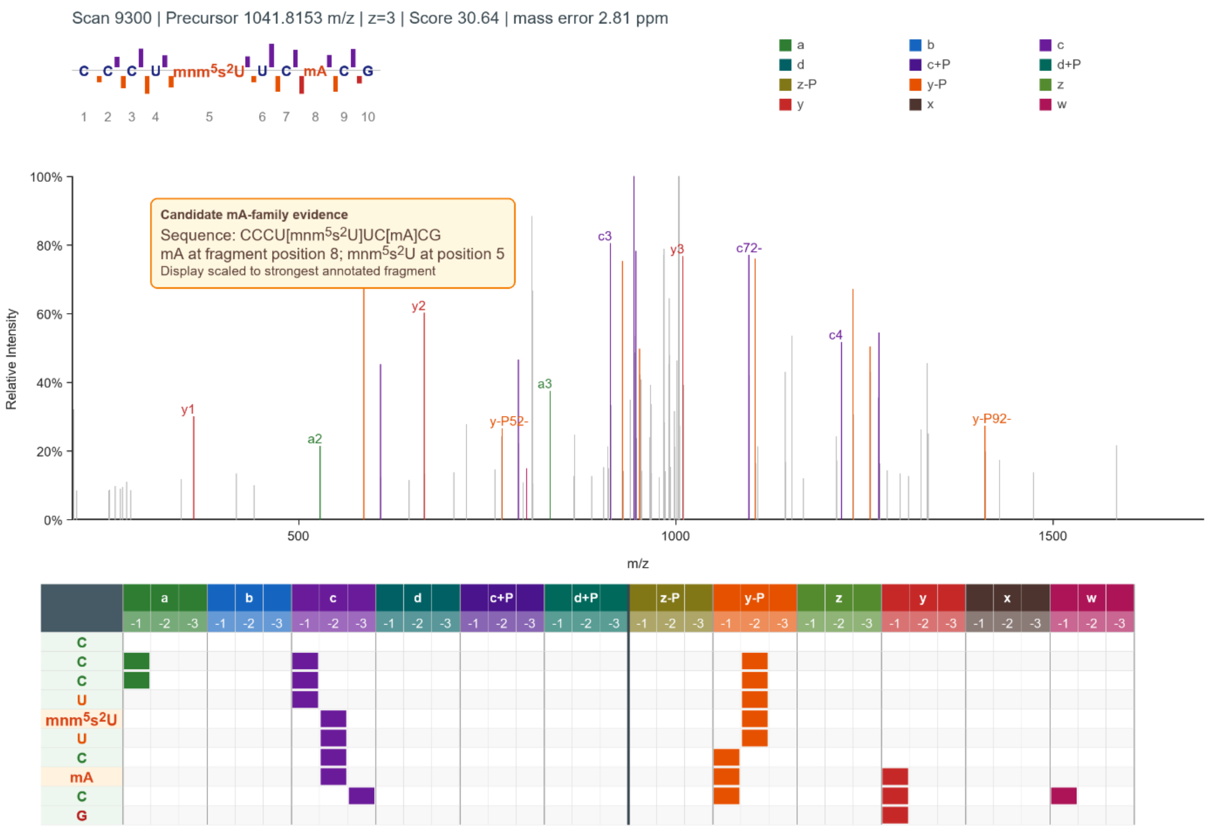


**Supplementary Figure 2. Annotated spectrum for a candidate mA-family signal at Sprinzl position 37 in *E. coli* tRNA.**

Annotated MS/MS spectrum generated from OligoFragger idXML annotation output and mzML peak data for scan 9300 from the RNase T1 E. coli tRNA dataset, assigned to CCCU[mnm5s2U]UC[mA]CG, illustrating fragment-ion support for a candidate mA-family signal in an anticodon-loop fragment that also contains mnm5s2U.

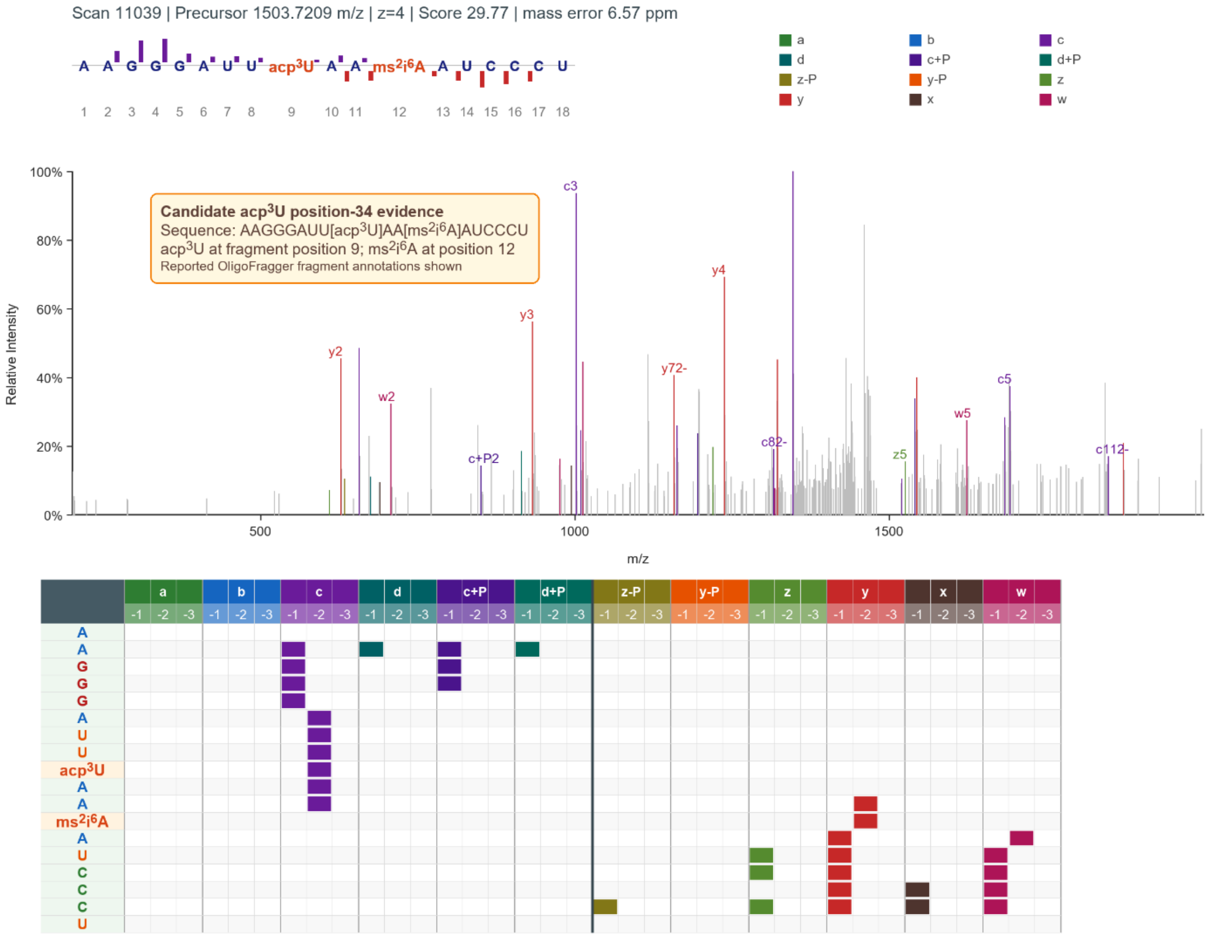


**Supplementary Figure 3. Annotated spectrum for a candidate acp3U signal at Sprinzl position 34 in *E. coli* tRNA.**

Annotated MS/MS spectrum generated from OligoFragger idXML annotation output and mzML peak data for scan 11039 from the RNase A E. coli tRNA dataset, assigned to AAGGGAUU[acp3U]AA[ms2i6A]AUCCCU, as representative spectrum support for the candidate acp3U signal discussed for Leu-UAA position 34.

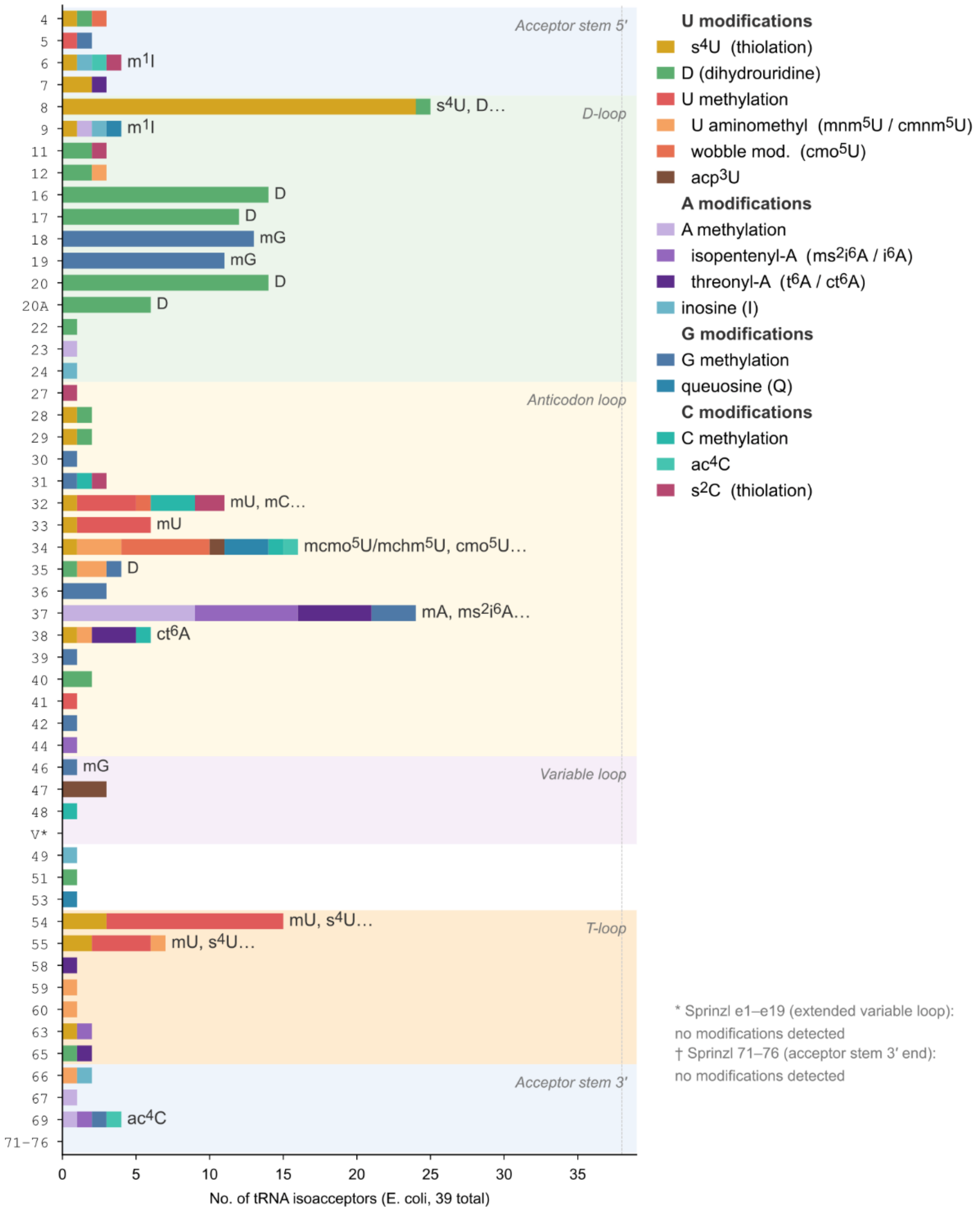


**Supplementary Figure 4.** ***E. coli*** **tRNA modification prevalence across Sprinzl positions.**

The chart summarizes the number of isoacceptors with non-suspicious modification calls at each Sprinzl position, grouped by modification category.

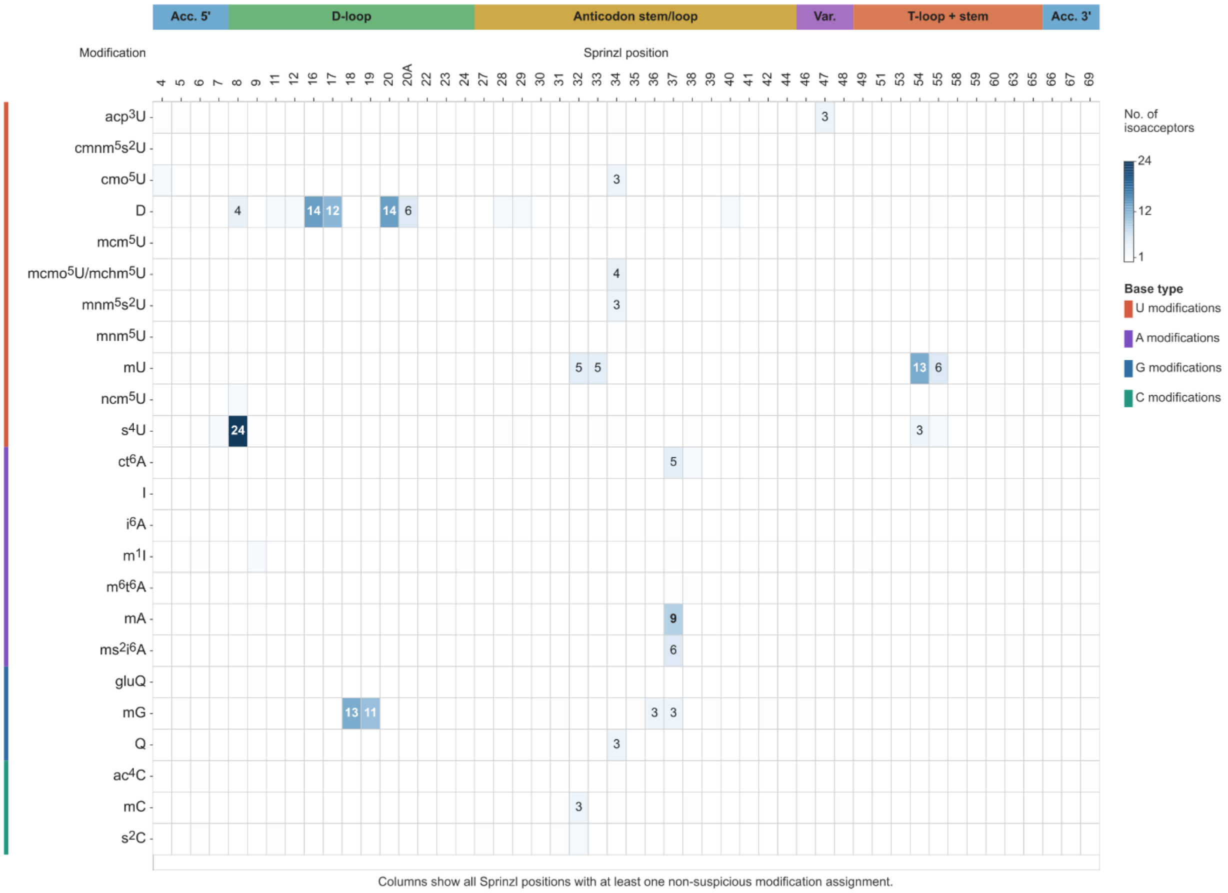


**Supplementary Figure 5.** ***E. coli*** **tRNA modification distribution across Sprinzl positions.**

Heatmap cells show the distribution of detected non-suspicious modification types across Sprinzl positions in the *E. coli* tRNA repertoire.

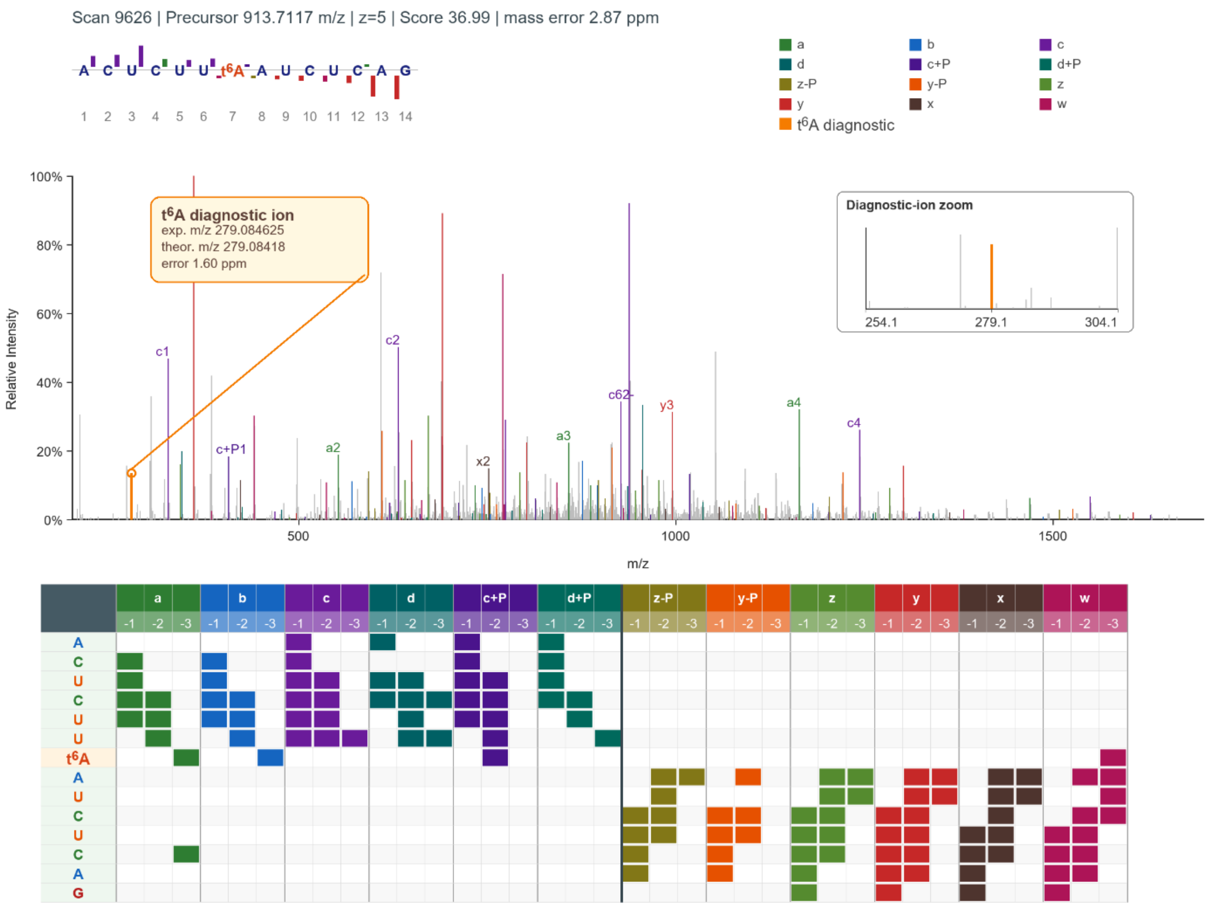


**Supplementary Figure 6. Representative annotated spectrum for a t6A-containing human tRNA fragment.**

Annotated MS/MS spectrum generated from OligoFragger idXML annotation output and mzML peak data for scan 9626 from HAP1_tRNA_clone1_rep1.mzML, displayed as the corrected sequence ACUCUU[t6A]AUCUCAG. The t6A diagnostic ion is annotated at experimental m/z 279.084625 (theoretical m/z 279.08418; 1.60 ppm error), together with fragment-ion evidence across the oligonucleotide.

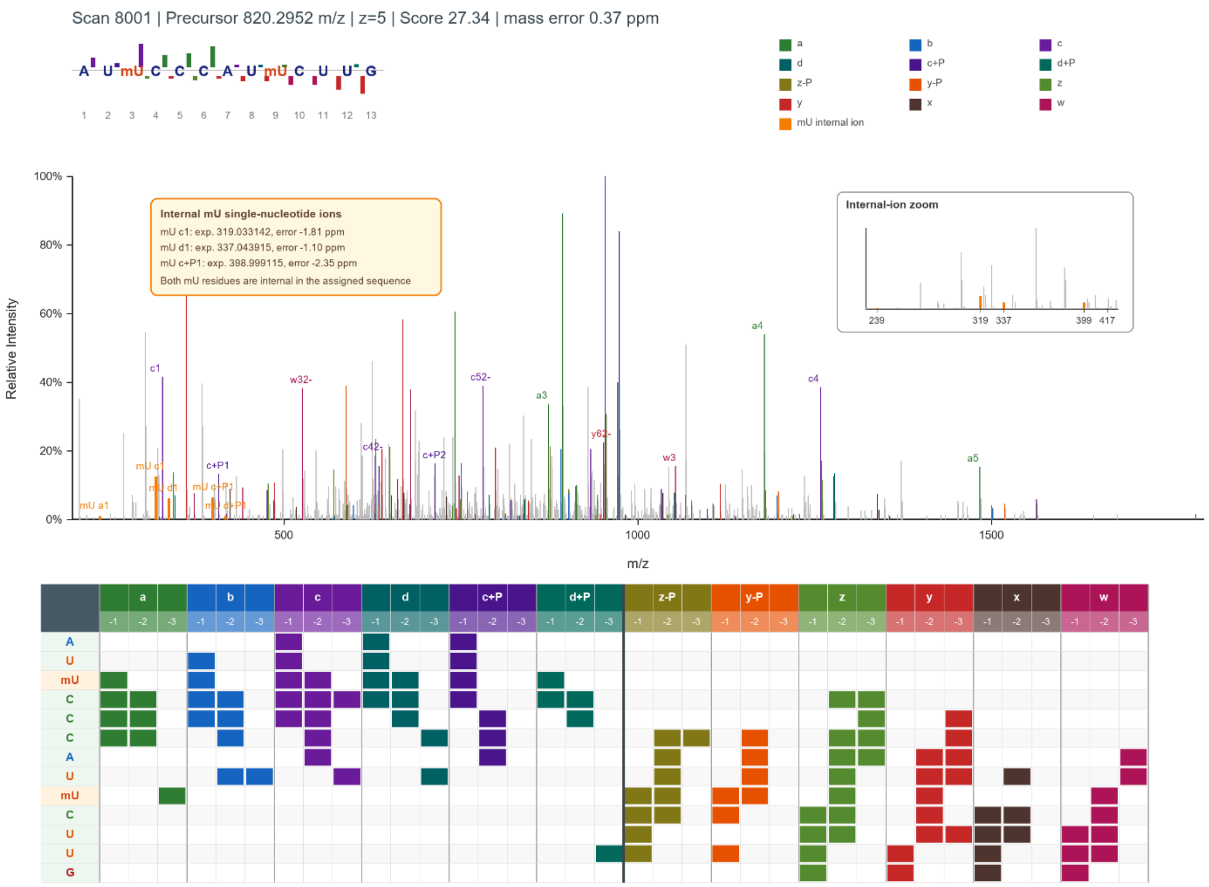


**Supplementary Figure 7. Representative annotated spectrum showing internal single-nucleotide evidence for a non-labile human tRNA modification.**

Annotated MS/MS spectrum generated from OligoFragger idXML annotation output and mzML peak data for scan 8001 from HAP1_tRNA_clone1_rep1.mzML, assigned to AU[mU]CCCAU[mU]CUUG. Although both mU residues are internal to the oligonucleotide, the spectrum contains mU-derived single-nucleotide ions at experimental m/z 319.033142 (mU c1; theoretical m/z 319.033719; -1.81 ppm error), 337.043915 (mU d1; theoretical m/z 337.044285; -1.10 ppm error) and 398.999115 (mU c+P1; theoretical m/z 399.000051; -2.35 ppm error).

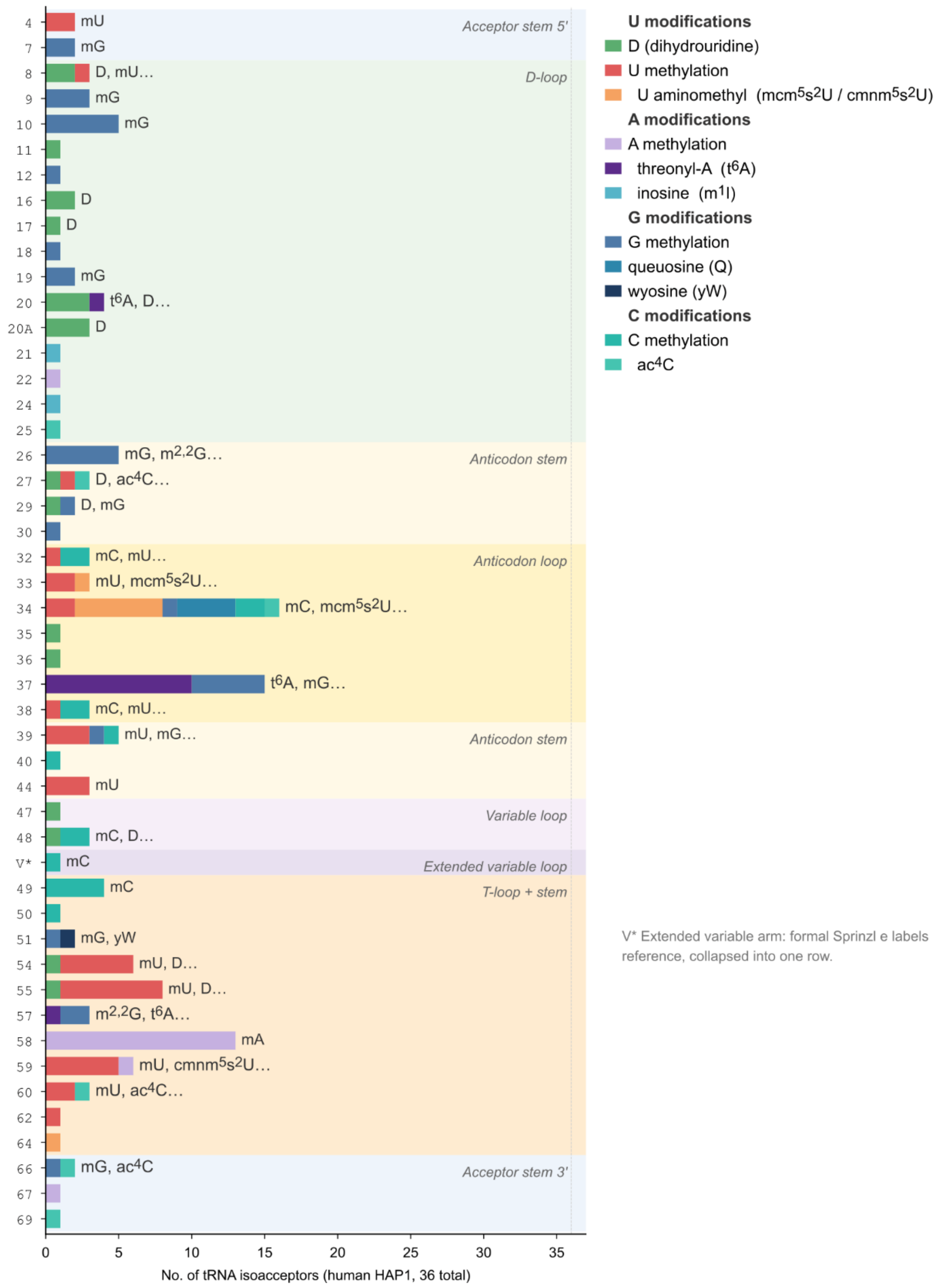


**Supplementary Figure 8. Human HAP1 tRNA modification prevalence by Sprinzl region.**

Each row represents a Sprinzl position or structural region and summarizes the number of isoacceptors carrying non-suspicious OligoFragger modification calls, grouped by modification pathway.

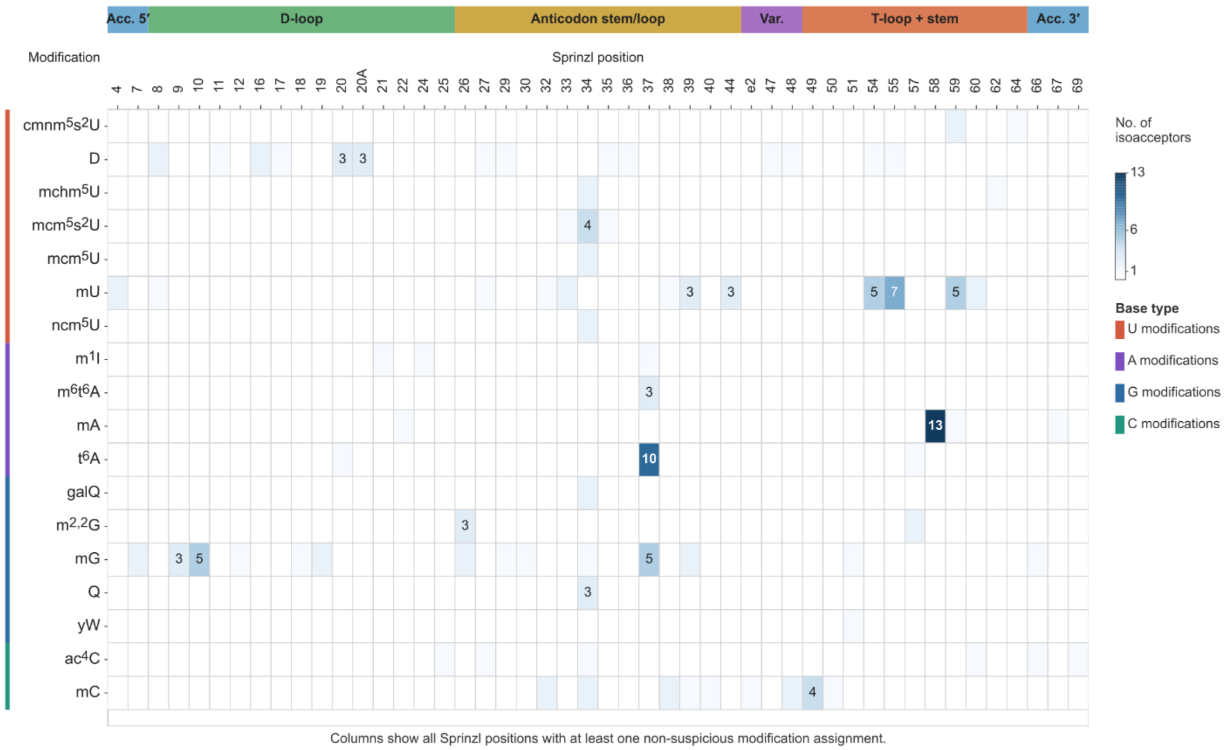


**Supplementary Figure 9. Human HAP1 tRNA modification distribution across Sprinzl positions.**

Heatmap cells show the number of isoacceptors supporting each modification type at each Sprinzl position after suspicious assignments were removed.

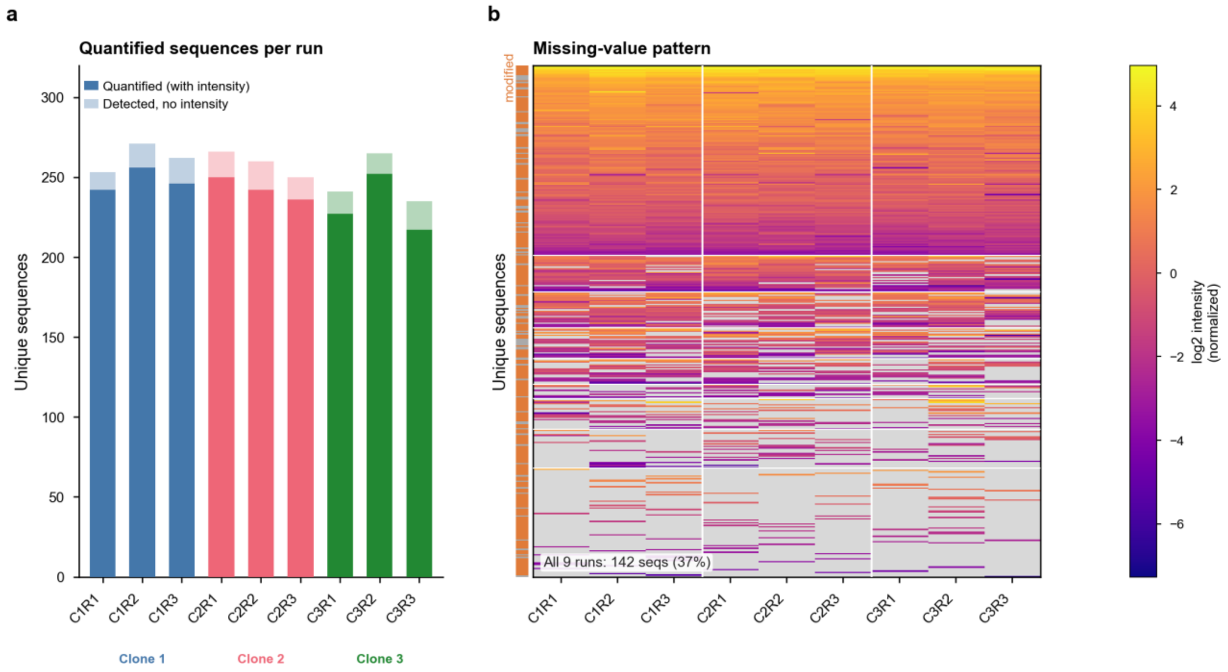


**Supplementary Figure 10. OSM-level quantification completeness for human HAP1 tRNA.** (a) Number of unique sequences quantified with IonQuant intensity versus detected but not quantified per run, color-coded by biological clone. (b) Missing-value heatmap for 382 unique quantified sequences across 9 runs, with colored cells indicating median-normalized log2 intensity and gray cells indicating missing values. Clone separators are shown as vertical lines, and the row annotation indicates modified or unmodified sequences. In total, 142 sequences (37%) were quantified in all 9 runs.

**Supplementary Notes**

**Supplementary Note 1. Terminal-state-aware precursor validation and ion-label convention.** RNA oligonucleotide MS/MS differs from peptide MS/MS in that the same nucleotide sequence can be observed with multiple 5' and 3' terminal states. RNase digestion and gas-phase fragmentation can generate fragments carrying 5'-OH or 5'-phosphate termini, together with 3'-OH, 3'-phosphate or dehydrated-phosphate terminal states, as reported in prior MS analyses of RNA cleavage products[1,2]. If these states were encoded as separate in silico digestion products, each sequence candidate would expand into multiple terminal variants before spectrum evidence was considered. OligoFragger instead keeps digestion products in nucleotide-sequence space and introduces terminal-state offsets during precursor validation.

OligoFragger follows the conventional RNA oligonucleotide fragmentation nomenclature in which a, b, c and d ions are fragments retaining the 5' side of the oligonucleotide, whereas w, x, y and z ions are fragments retaining the 3' side[3-5]. These labels are used in manuscript spectra and OligoFragger annotation output to describe the chemical termini of observed low-mass and backbone-fragment ions. Because RNA oligonucleotides can carry different terminal phosphate states, OligoFragger uses an explicit display convention that preserves the familiar ion names while indicating whether a terminal phosphate is present or absent relative to the reference state.

The reference annotation state used for display is an RNA fragment carrying a 3'-phosphate terminus. Under this convention, standard a, b, c, d, w, x, y and z retain their usual RNA-fragment interpretation. Additional labels are used only to make terminal-state evidence explicit: c+P and d+P denote 5'-side ions carrying an additional phosphate relative to the corresponding c or d labels, whereas z-P and y-P denote 3'-side ions lacking the phosphate present in the corresponding z or y reference labels. These +P and -P suffixes should be read as annotation labels for terminal-state evidence rather than as separate fragmentation mechanisms.

| OligoFragger label group | Terminal evidence represented | Use in this manuscript |
|---|---|---|
| **a, b, c, d** | 5'-side ions without an added 5' phosphate | Conventional prefix-side sequence evidence |
| **c+P, d+P** | 5'-side ions with an added phosphate | Evidence for a 5'-phosphate-containing terminal state |
| **z-P, y-P** | 3'-side ions lacking the reference 3' phosphate | Evidence for a 3'-OH terminal state |
| **z, y** | 3'-side ions in the 3'-phosphate reference convention | Evidence for a dehydrated-phosphate-associated 3' terminal state in precursor-offset validation |
| **x, w** | 3'-side ions retaining phosphate-containing termini | Evidence for a 3'-phosphate-containing terminal state |

The same observed mass can sometimes be compatible with complementary RNA ion interpretations under a fixed terminal state. OligoFragger therefore uses these labels as spectrum annotations and terminal-state evidence after route matching, while final prefix or suffix assignment is determined by compatibility with the candidate sequence route rather than by the low-mass ion label alone.

The terminal offsets evaluated in OligoFragger are no offset, HPO3, HPO3-H2O, 2HPO3 and 2HPO3-H2O. After spectrum-graph routes are matched to a candidate sequence, the terminal evidence classes above are used to restrict the terminal offset applied during precursor validation when both sides are informative.

| 5' terminal evidence | 3' terminal evidence | Precursor offset |
|---|---|---|
| **5'-OH** | 3'-OH | no offset |
| **5'-OH** | 3'-dehydrated phosphate | HPO3-H2O |
| **5'-OH** | 3'-phosphate | HPO3 |
| **5'-phosphate** | 3'-OH | HPO3 |
| **5'-phosphate** | 3'-dehydrated phosphate | 2HPO3-H2O |
| **5'-phosphate** | 3'-phosphate | 2HPO3 |

When terminal evidence is incomplete or contradictory, OligoFragger does not force a single terminal-state assignment; instead, the configured terminal-offset search remains available during precursor validation. This design avoids assuming a fixed terminal phosphate state during database digestion while still allowing spectrum evidence to reduce the number of precursor-compatible terminal states.

**Supplementary Note 2. Suspicious modification annotation.** OligoFragger reports suspicious modification assignments as site-level spectrum-evidence annotations. These annotations are not used to remove OSMs during target-decoy competition or q-value calculation. They are intended to identify modification sites for which the accepted sequence assignment remains reported, but the modification at a specific residue is less strongly supported than a simpler spectrum-supported explanation.

For internal modified positions, OligoFragger evaluates the selected prefix and suffix routes that support the accepted OSM. A modified residue is compared against alternative route candidates that cover the same absolute sequence position, preserve the already matched local sequence context from that route direction and propose a less-modified residue or composition at the tested position. If the less-modified competing route has stronger mass-accuracy-weighted intensity support than the route used in the accepted assignment, the modified site is marked as suspicious from that route-side evidence. This comparison is therefore based on the observed spectrum and matched route evidence, not on whether the modification would be chemically plausible in a particular RNA type.

This route-level comparison is distinct from final OSM selection. OligoFragger does not choose the best local route alone as the final assignment: the final OSM must also explain the observed precursor mass after isotope and terminal-state offsets are considered. Therefore, a less-modified route can have stronger local fragment evidence but not be selected in the original search because removing or reducing the modification would make the candidate mass incompatible with the precursor. In this situation, the less-modified route is not a valid final candidate, but it still indicates that the local evidence for the modified residue is weak. The suspicious flag therefore does not replace the accepted OSM with the less-modified route; it records that the modification at that residue is weakly supported relative to a simpler local spectrum-supported explanation.

Terminal positions require a separate check because the route-start evidence can be represented by one- or two-nucleotide graph nodes. For terminal modified dinucleotides, OligoFragger compares matched two-nucleotide ion evidence with the same unmodified dinucleotide composition. If a less-modified terminal composition has stronger total intensity support than the modified composition used by the accepted route, the modified terminal residue or residues are marked as suspicious. This rule is designed to avoid over-interpreting terminal low-mass evidence when a simpler composition explains the same local sequence context better.

Inosine ([I]) is handled especially conservatively because inosine/deamination-related evidence can be prone to false assignment, including near-isotope-offset alternatives to adenosine-containing evidence. An [I] assignment is therefore marked suspicious unless the position is covered by both prefix- and suffix-side route evidence. This additional rule is specific to [I] and should not be interpreted as a general biochemical plausibility filter.

A suspicious modification flag does not mean that the OSM is false, that the residue is chemically impossible or that the modification site is merely localization-ambiguous. It means that, within the accepted OSM, the spectrum contains stronger or insufficiently corroborated evidence for a less-modified explanation at that site. For biological summaries in this manuscript, suspicious modification sites were excluded, whereas the corresponding OSMs remained part of the spectrum-level search output and FDR-controlled identification set.
